# PanGene-O-Meter: Intra-Species Diversity Based on Gene-Content

**DOI:** 10.1101/2025.09.01.673544

**Authors:** Haim Ashkenazy, Detlef Weigel

## Abstract

**Background:** Bacterial genome evolution is shaped extensively by horizontal gene transfer, generating gene presence-absence variation that represents a major axis of functional diversity yet is largely invisible to core-genome-based similarity metrics. While genome-wide average nucleotide identity (ANI) remains the most widely used measure of similarity, it considers only the alignable fraction of compared genomes and therefore fails to capture gene-content differences that can have profound phenotypic consequences even among nearly identical strains.

**Results:** Here we introduce PanGene-O-Meter, a computational framework that quantifies gene-content similarity (GCS) between bacterial genomes by leveraging pan-genome structure and orthology group relationships. Applying PanGene-O-Meter to four large datasets spanning *Escherichia coli*, *Pseudomonas aeruginosa*, *Staphylococcus aureus*, and *Helicobacter pylori*, we show that GCS captures substantial functional diversity hidden within high-ANI genome pairs, and that gene-content-based trees resolve substructure within sequence types that core-genome analyses cannot. We further introduce GeneContRep, an efficient greedy algorithm for gene-content-based genome dereplication, and validate its utility on 1,000 *Klebsiella pneumoniae* genomes from two major multidrug-resistant lineages (ST258 and ST307) with matched experimental antimicrobial resistance phenotypes. GeneContRep preserves substantially more clinically relevant diversity than standard ANI-based dereplication, recovering carbapenemase variants, capsule locus types, and resistance determinants that are systematically lost under conventional ANI thresholds.

**Conclusions:** PanGene-O-Meter provides an efficient, complementary, and biologically interpretable framework for quantifying intra-species bacterial diversity, with broad applications in comparative genomics, epidemiological surveillance, and the principled selection of representative genomes for functional studies. PanGene-O-Meter is available at: https://github.com/HaimAshk/PanGene-O-Meter.

## Introduction

The classification and grouping of bacteria into taxonomic and functional units has been a long-standing and often intensely debated challenge (Rosselló-Mora 2005; Konstantinidis et al. 2006). Clustering of bacterial strains by different measures of similarity has significant practical implications, including tracking pathogenic outbreaks, monitoring antibiotic resistance, and understanding the structure and evolution of microbial communities (reviewed by Van Rossum et al. 2020). The availability of fully sequenced bacterial genomes and metagenomes has revealed not only extreme intra-species diversity of bacteria, but also their extensive evolutionary plasticity. This shift has led to a transition from morphological and biochemical-based classification schemes to genomic marker-based and whole-genome-based approaches (Richter and Rosselló-Móra 2009; Ludwig and Klenk 2001). The rapid growth in the availability of complete genomes from individual isolates and, more recently, metagenome-assembled genomes (MAGs) has also brought up a related problem: What is the best strategy for identifying the optimal representative genome of a taxonomic unit for further functional studies, especially when dealing with fragmented or incomplete assemblies?

Average nucleotide identity (ANI) is currently the most widely used metric for whole-genome-based quantification of bacterial similarity. ANI is based on the pairwise alignment of bacterial genomes, quantifying their identity in the alignable DNA sequence space. Numerous variants and efficient implementations of ANI calculation have been proposed, including focusing on coding regions, reciprocally homologous regions, and applying different sequence alignment methods (Goris et al. 2007; Richter and Rosselló-Móra 2009; Pritchard et al. 2016; Jain et al. 2018; Shaw and Yu 2023; Varghese et al. 2015). Fast estimation of ANI can be achieved by employing kmer-based methods, such as calculation of MinHash distance, as implemented in MASH (Ondov et al. 2016) and sourmash (Irber et al. 2022).

Even within designated bacterial species, gene content can differ substantially (Nysten et al. 2024), primarily resulting from rapid gene acquisition through horizontal gene transfer (HGT) (Ochman et al. 2000; Koonin and Wolf 2008), integration of other mobile elements (e.g., prophages, and integrative conjugating elements), as well as rampant gene loss events (Bolotin and Hershberg 2015; Snel et al. 2002). Such gene-content variation can have significant phenotypic and functional impact (Kieffer et al. 2025; Adams et al. 2025). Notable examples include pathogenicity islands and antibiotic resistance genes, which are major factors in determining host-range of pathogenic bacteria (Hacker and Carniel 2001; Dobrindt et al. 2004). More recently, the discovery of biosynthetic pathways (Gavriilidou et al. 2022) as well as numerous anti-phage defense systems that individually are often rare (Labrie et al. 2010; Doron et al. 2018; Hussain et al. 2021; Hochhauser et al. 2023; Millman et al. 2022; Getz et al. 2025) have contributed to our understanding of the functional diversity imposed by such gene-content variation. For example, the presence or absence of specific defense genes generates substantial phenotypic differences between bacterial strains that have otherwise very similar genomes, with high ANI values (Hussain et al. 2021).

A drawback of the existing measures, such as ANI, is their limited ability to capture gene-content diversity, since they consider only the homologous (alignable) parts of the analyzed genomes. In other words, regions not present in a subset of the analyzed genomes are not directly accounted for. Because ANI does correlate, at least to some extent, with gene-content, different cutoffs of ANI values have been suggested to be useful when assigning membership to groups of bacterial strains, often referred to as the ‘ANI gap’ (Rodriguez-R et al. 2024; Viver et al. 2024), but it can by design be only a proxy for gene-content similarity (Van Rossum et al. 2020).

An alternative to the ANI scheme for strain tracking was recently suggested, making use of genome synteny (Enav et al. 2024). The SynTracker method quantifies similarity between bacterial genomes based on the order of sequence blocks in homologous genomic regions. By focusing on structural changes in the genomes rather than point mutations, SynTracker can provide an additional layer of information for the classification of bacterial genomes. However, SynTracker, like ANI, relies on homologous regions and does not directly account for genes missing in some of the analyzed genomes.

Unlike ANI and SynTracker, which are confined to homologous regions, PopPUNK does capture accessory-genome divergence directly, estimating core and accessory distances through a fast k-mer-based decomposition (Lees et al. 2019). While highly scalable and well suited to broad epidemiological tracking, its reliance on k-mer distributions still precludes the direct biological identification of the specific genes driving that divergence.

Resolving which genes drive this divergence requires an explicit, gene-level representation of genome content, precisely what the pan-genome concept provides. The pan-genome represents the total gene-content within all the known genomes of a bacterial taxon (Lapierre and Gogarten 2009; Tettelin et al. 2008, 2005; Hogg et al. 2007; Dewar et al. 2024). Groups of orthologous genes within a pan-genome are termed pangenes, and depending on their frequency across the analyzed genomes, they are assigned to the “core genome,” the “accessory” or “variable genome,” or the “shell genome.” Each bacterial genome can thus be represented by the presence-absence profile of its pangenes, often termed a ‘phyletic pattern.’ Because it records gene content at the level of individual pangenes, the phyletic pattern makes it possible to visualize and compare gene-content across genomes, to identify gene gain and loss, to discern functional dependencies between genes, and to reconstruct the evolutionary history of bacterial lineages (Tettelin et al. 2005; Cohen and Pupko 2010; Cohen et al. 2013). This same gene-level resolution provides a direct basis for comparing genomes by functional content, and thus for identifying a functionally representative genome.

Here, we introduce PanGene-O-Meter, a method that employs phyletic patterns to directly quantify the gene-content similarity (GCS) among a given set of bacterial genomes. We demonstrate its usefulness by analyzing bacterial genomes of four bacterial species: the gram-negative bacteria *Escherichia coli*, *Helicobacter pylori* and *Pseudomonas aeruginosa*, and the gram-positive species *Staphylococcus aureus*. We demonstrate that core-genome identity decouples substantially from accessory gene dynamics among closely related strains (ANI > 99.5%). Furthermore, we provide an efficient method for accurate GCS estimation using the fast protein clustering approach implemented in DIAMOND (Buchfink et al. 2026). Finally, we apply PanGene-O-Meter and our greedy clustering algorithm, GeneContRep, to dereplicate a curated clinical cohort of 1,000 multidrug-resistant *Klebsiella pneumoniae* genomes. We demonstrate that gene-content-based dereplication preserves critical clinical diversity, including antimicrobial resistance determinants and capsule types, that is otherwise erased by standard ANI-based dereplication. All together, PanGene-O-Meter provides a highly interpretable and accurate method for facilitating the selection of ‘representative’ genomes for downstream functional genomics.

## Results

### Genome quality and gene-content variability across the four species datasets

We start by characterizing genome assemblies of isolates belonging to four distinct bacterial species: *Escherichia coli* (700 genomes, 50 for each phylogroup (Abram et al. 2021)), *Helicobacter pylori* (500 genomes), *Pseudomonas aeruginosa* (500 genomes), and *Staphylococcus aureus* (500 genomes). Previous studies have demonstrated that these species, which are well-represented in genomic databases, are characterized by different HGT rates and therefore are expected to show different levels of gene-content diversity (Oliveira et al. 2017). In addition, *P. aeruginosa* and *S. aureus* are of major public health relevance and are part of the 2024 WHO Bacterial Priority Pathogens List (World Health Organization 2024). Only a few of the genomes were de novo annotated; for the majority, NCBI gene annotations were used (see methods).

As expected, based on their HGT rates, the variance in total genome size and the number of genes per genome is highest among *P. aeruginosa* isolates, followed closely by *E. coli*, with an intermediate level in *S. aureus*, and the lowest for *H. pylori* (Supplementary Figure S1 and Supplementary Table S1). Importantly, this variability cannot be attributed to the quality and completeness of the genome assemblies, as reflected by the low intra-species variance of these measures in all species (Supplementary Figure S1 and Supplementary Table S1). The reconstructed species pan-genomes also reflect this variability, with *E. coli* having the highest number of gene clusters and the highest number of accessory genes (Supplementary Figure S1E); we note that *E. coli* was sampled more deeply and across all phylogroups (700 vs. 500 genomes), which contributes to its larger pan-genome, so this cross-species comparison reflects both intrinsic plasticity and sampling design. We further identified the sequence types (STs) for each genome using the PubMLST scheme. STs are determined by specific combinations of alleles at multiple conserved loci through a methodology known as Multilocus Sequence Typing (MLST). The PubMLST database functions as a curated repository, offering standardized typing schemes for diverse microbial species (Jolley et al. 2018). The same trend of variability was observed with respect to the number of predicted STs (Supplementary Table S1).

### A new method for quantifying genome similarity based on gene content

To go beyond average nucleotide identity in the shared sequence space and to also account for relationships based on genes with presence/absence patterns across the studied groups of genomes, we implemented the following steps:

1. **Protein-coding gene prediction:** Identify protein-coding genes in each genome assembly (usually already provided through the available genome annotation).
2. **Orthology group assignment:** Assign predicted genes to orthology groups (pangenes) to construct a pan-genome (different methods can be used for this step).
3. **Phyletic pattern representation**: Represent each genome as a binary presence/absence (P/A) vector, where each element corresponds to a pangene and the value is either 1 (present) or 0 (absent).
4. **Gene-content similarity (GCS) calculation**: Calculate GCS based on Jaccard similarity (“GCSj”) (Eq. 1) between the P/A vectors of two bacterial genomes (*pS*1 and *pS*2):

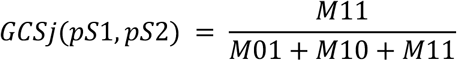

where *M*11 represents the total number of elements (orthology-groups) for which both *pS*1 and *pS*2 have a value of 1; *M*01 represents the total number of elements for which the value of *pS*1 is 0 and *pS*2 is 1; and *M*10 represents the total number of elements for which the value of *pS*1 is 1 and the value of *pS*2 is 0.

Alternatively, the GCS can be defined by the shared pangenes, based on their overlap coefficient (“GCSo”). This approach reduces the impact of partial genomes on the GCS metric (Eq. 2):

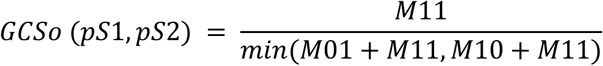

For each of the analyzed datasets, PanGene-O-Meter uses PanX (Ding et al. 2018) to assign protein sequences into orthology groups, which serve as the basis for computing the GCS measures as described above.

We first compared the performance of several similarity measures: ANI, GCSj, GCSo, Average Pairwise Synteny Scores (SynTracker), and PopPUNK accessory similarity (Figure 1A and Supplementary Figures S2-S4). As expected, core-genome-based metrics ANI and SynTracker are highly correlated across all datasets (Pearson correlation coefficients [PCCs] ranging from 0.817 to 0.963). However, this correlation drops substantially in the more diverse species when comparing very closely related strains with ANI values between 99 and 100 (PCCs of 0.47, 0.35, 0.94, and 0.84 for *E. coli*, *P. aeruginosa*, *S. aureus*, and *H. pylori*, respectively).

**Figure 1.**
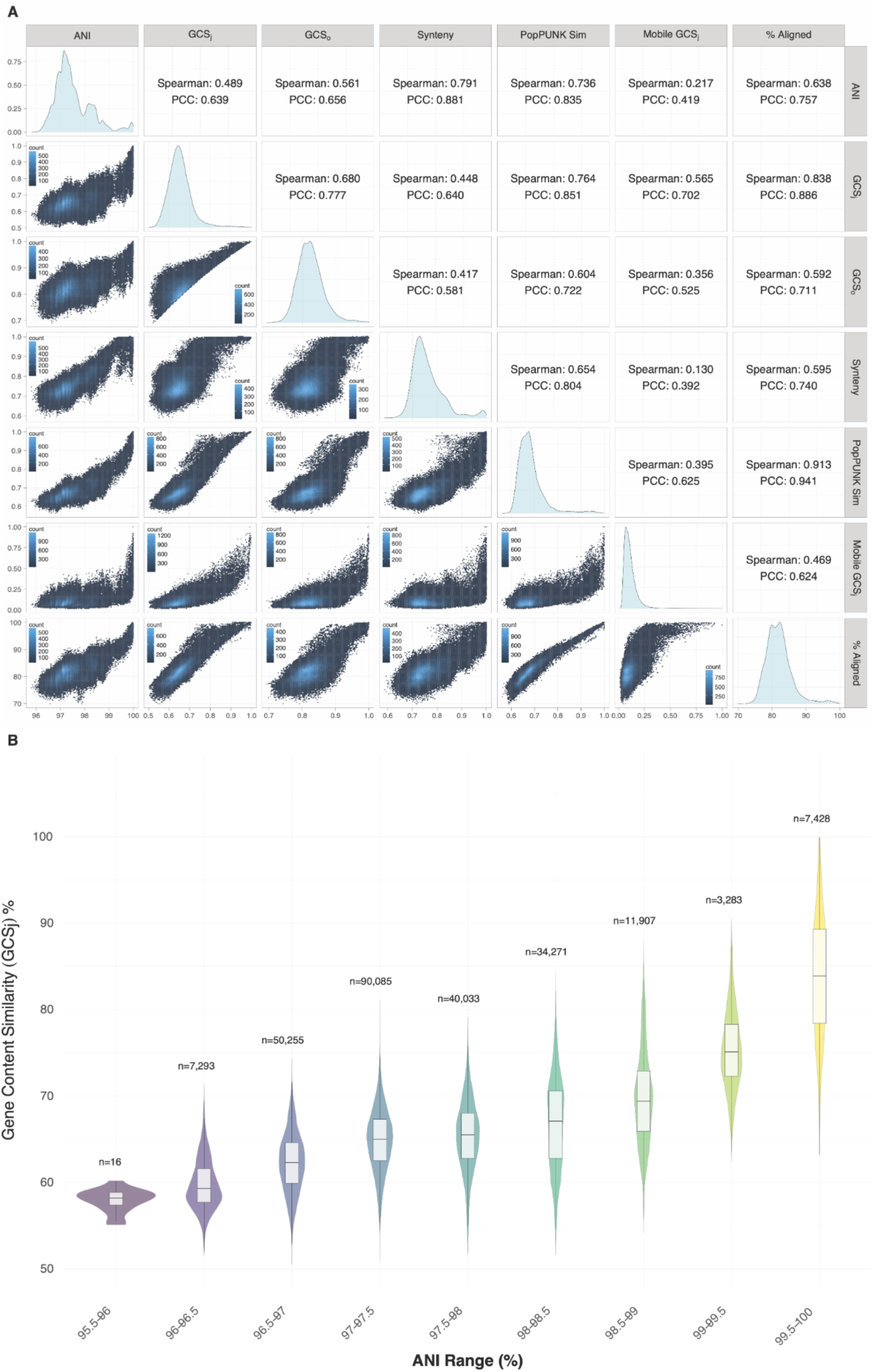
Comparing pairwise similarity measures for *E. coli* genomes. **(A)** Pairwise comparisons of similarity measures: ANI; gene-content similarity by Jaccard (GCSj) and overlap coefficient (GCSo); synteny score (SynTracker); PopPUNK accessory similarity; mobile-element profile similarity (Mobile GCSj); and fraction of aligned bases (% Aligned). Upper-right panels give Pearson (PCC) and Spearman coefficients. Gene-content similarity varies widely even among pairs with near-identical ANI, as quantified in (B). No points fall below the diagonal in the GCSj–GCSo panel because GCSo ≥ GCSj by definition (Eq. 2). **(B)** Distribution of GCSj values across binned ANI ranges. Violin plots show the density distribution of GCSj, with internal boxplots indicating the median and interquartile range (IQR). The number of pairwise comparisons (*n*) is shown above each bin. The variance of GCSj is not homogeneous across ANI ranges (Fligner-Killeen test, p-value < 2.2e-16), and the spread of gene-content similarity peaks at the highest sequence identities (ANI 99.5-100%), highlighting extensive accessory genome variation among closely related strains.

To evaluate accessory-genome-based metrics, we compared GCSj against PopPUNK, a well-established k-mer-based method for estimating core and accessory distances (Lees et al. 2019). We observed a strong correlation between PopPUNK’s accessory similarity and GCSj across most datasets (PCC = 0.878, 0.890, and 0.880 for *E. coli*, *P. aeruginosa*, and *H. pylori*, respectively; a more moderate 0.651 for *S. aureus*), providing external validation that both methods capture similar biological signals regarding accessory genome dynamics. Because PopPUNK estimates these distances only from k-mer distributions, additional analyses would be required for biological interpretation. In contrast, PanGene-O-Meter explicitly exploits orthology group assignments, allowing for direct identification of the specific genes and mobile elements driving strain divergence.

The differences between ANI and the gene-content-based measures (GCSj and GCSo) become even more pronounced among closely related strains. To quantify this decoupling of sequence similarity and gene content, we evaluated the variance of GCSj within binned ANI ranges (Figure 1B and Supplementary Figures S2-S4). We observed significant heteroscedasticity across ANI bins in all four species (Fligner-Killeen test, p-value < 2.2e-16). Notably, in *E. coli*, *P. aeruginosa*, and *S. aureus*, the spread of GCSj values reaches its maximum among the most closely related strains (ANI 99.5-100%), with interquartile ranges (IQR) of 9.7%, 7.7%, and 5.4%, respectively. This indicates that even when the core genome is nearly identical, strains can exhibit extensive accessory genome variation. The exception is *H. pylori*, which consistently displays low gene-content variance across all populated ANI ranges (IQR ≤ 3.4% throughout, and only 0.5% at ANI > 99.5%), in agreement with its limited accessory genome diversity (Supplementary Figure S1E).

The correlation between GCSj and GCSo is relatively high across all datasets (PCCs ranging from 0.72 to 0.87; Figure 1A and Supplementary Figures S2-S4). By design, GCSo calculates similarity based solely on the size of the smaller genome (the overlap coefficient). This approach is expected to be more robust when comparing genome assemblies with high degrees of incompleteness, as it reduces the penalizing effect of ‘missing’ genes that are merely artifacts of the sequencing or assembly process. To empirically validate this, we simulated random losses of genes from the genome sequence (assembly incompleteness) in the *E. coli* phyletic-pattern matrix and tracked the resulting error in our similarity metrics (Supplementary Figure S5). When incompleteness was simulated in both genomes of a pair simultaneously, GCSj and GCSo performed comparably, with both showing modest increases in error. The difference became striking when only one genome assembly in a pair was degraded, a scenario mimicking the comparison of a highly complete reference genome against an incomplete metagenome-assembled genome (MAG). In this case, GCSo remained highly robust, maintaining a near-zero median error even at 30% simulated incompleteness, whereas GCSj error increased linearly with the degree of incompleteness. Given the high completeness of the assemblies analyzed here (median >99% for all species; Supplementary Table S1), GCSj is the primary metric used throughout this study. However, we recommend GCSo for datasets containing MAGs or other incomplete assemblies.

Ultimately, the most important consideration is whether the observed gene-content variation is biologically relevant rather than an artifact of assembly or gene calling. To this end, we correlated GCSj and GCSo with the fraction of aligned bases. As expected, there is a strong correlation between these values in all datasets, indicating that the more unaligned bases, the lower the pairwise gene-content similarity. Furthermore, we annotated ‘mobile’ elements, including prophages, plasmids, genomic islands, antimicrobial resistance determinants, stress response genes, and virulence genes in each of the genomes and calculated the Jaccard similarity between profiles of these elements (Fig. 1 and Supplementary Figures S2-S4). Across datasets, the GCS measures correlated with mobile-element content comparably to PopPUNK (PCCs reported in Figure 1 and Supplementary Figures S2-S4), supporting the hypothesis that a large portion of the gene-content diversity can be attributed to variation of these elements, which can have a crucial effect on the biological activity spectrum of the studied taxa. Taken together, these results demonstrate that gene-content measures are highly effective at capturing functional and biological differences among closely related strains that would otherwise be obscured by high ANI values.

To visualize the relationship between core-genome and gene-content similarity in a common genome ordering, we generated nested ANI-GCSj heatmaps (Supplementary Figures S6-S9; Methods). Genomes were first grouped into coarse ANI-based clusters, and within each ANI group, genomes were reordered by GCSj similarity; the same final ordering was applied to both matrices, allowing direct comparison of nucleotide- and gene-content-based similarity structure. The resulting heatmaps show that ANI and GCSj share a broad similarity structure, consistent with the expected relationship between core-genome relatedness and gene-content similarity. Within ANI-coherent groups, however, GCSj revealed additional internal sub-structure, visible as finer banding and sub-block patterns not resolved by ANI alone, particularly in *E. coli*, *P. aeruginosa*, and *S. aureus*. This suggests that GCS can provide higher resolution in grouping bacterial genomes. *H. pylori* showed comparatively little internal GCSj sub-structure, consistent with its lower gene-content variability described above.

We next assessed how ANI and GCSj relate to maximum-likelihood (ML) distances computed from (i) a concatenated core-genome nucleotide alignment and (ii) the binary phyletic pattern used to compute GCSj (Supplementary Figure S10). ML distances from the core-genome alignment correlated strongly with ANI (PCCs -0.93 to -0.99). The correlation between GCSj and ML distances from the phyletic pattern was comparable (PCCs -0.96 to -0.98). In contrast, core-nucleotide ML distances and gene-content (phyletic-pattern) ML distances were only weakly to moderately correlated (PCCs +0.30 to +0.63), indicating that sequence divergence and gene-content divergence capture partially distinct signals. These results suggest that GCSj provides a good proxy for gene-content diversity and captures dimensions of variation that are not reflected by core-genome divergence (as measured by ANI or core-genome ML distances).

We next compared ML phylogenetic trees inferred from the core-genome alignment and from the full phyletic pattern across the four datasets (Fig. 2). In both tree types, isolates were largely grouped by STs, though ST clusters are somewhat less sharply delineated in the gene-content trees. The gene-content tree also resolved additional substructure within some STs. For example, in *E. coli* the gene-content tree split ST245 into two subgroups (Fig. 2A); this subdivision is driven by a set of orthologous genes present in subgroup 245.1 but absent from 245.2 (Supplementary Fig. S11).

**Figure 2:**
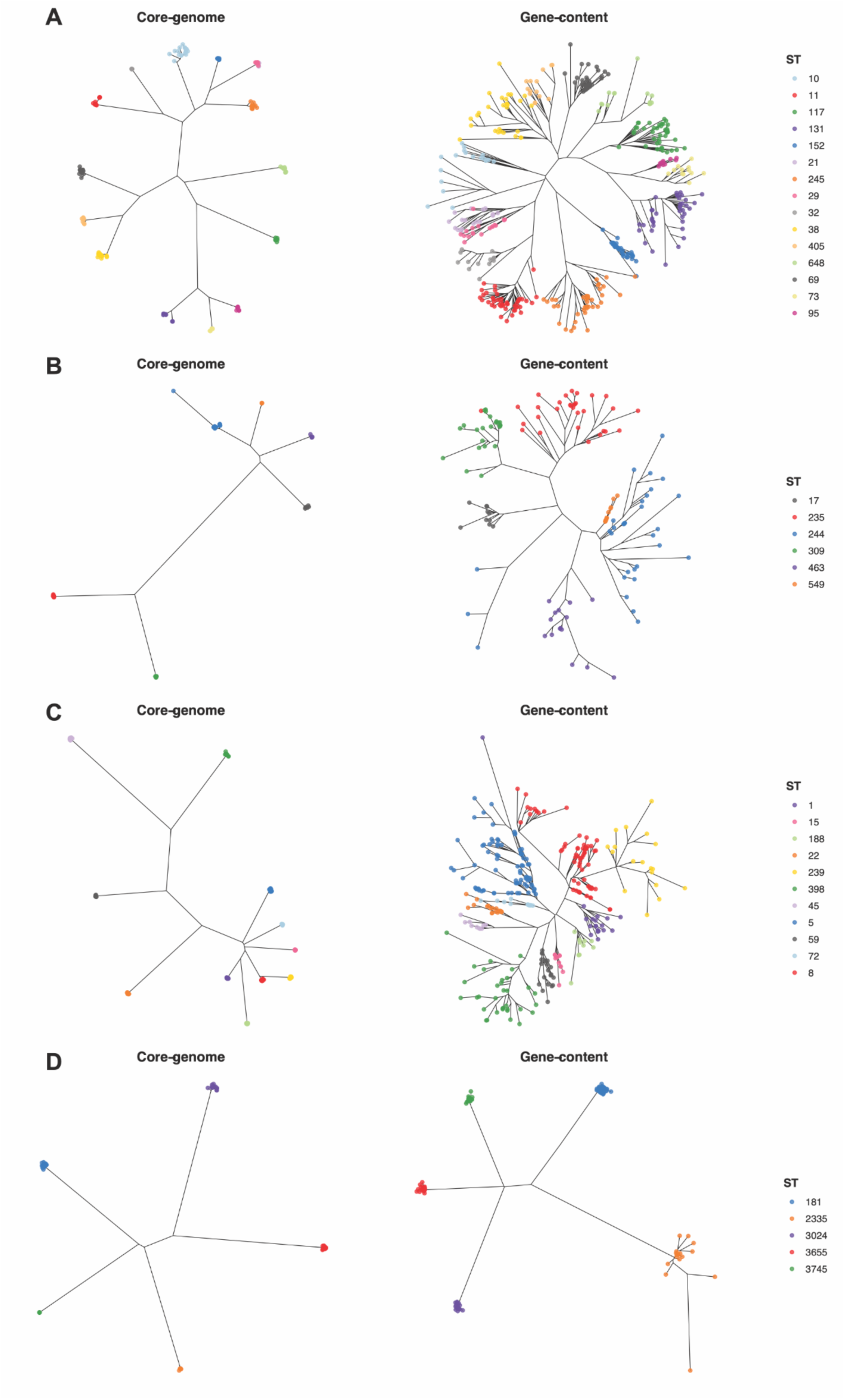
Maximum likelihood trees based on core-genome and gene-content for (A) *E. coli*, (B) *P. aeruginosa,* (C) *S. aureus*, and (D) *H. pylori*. Isolates are colored by their sequence type (ST). Only strains belonging to STs with at least 10 isolates were included in the analysis. Gene-content-based trees can reveal additional substructure within some STs.

Overall, grouping genomes by gene content (e.g., using GCSj) can yield a refined sub-classification beyond that provided by MLST or core-genome/ANI-based analyses, as illustrated by the ST245 subdivision. Gene content thus offers a complementary axis of variation that can reveal functionally relevant structure not captured by substitution-based trees, with its usefulness depending on the taxa and questions at hand. Gene-content trees should not be interpreted as phylogenies in the strict sense: gene presence/absence is shaped substantially by horizontal transfer and convergent loss, so these trees summarize combined vertical and horizontal signal rather than estimating vertical descent.

### Fast Estimation of Gene-Content Similarity

Accurate assignment of genes to orthology groups is a computationally intensive task that naively scales quadratically with the number of genomes (Ding et al. 2018). Recent advances in protein sequence clustering algorithms have enabled efficient clustering of millions and even billions of sequences (Steinegger and Söding 2017; Buchfink et al. 2026). We explored the applicability of these algorithms for fast estimation of gene-content similarity. Specifically, we employed the DIAMOND DeepClust bi-directional coverage clustering algorithm to cluster all proteins from all genomes for each of the analyzed datasets. The resulting clusters were treated as orthology groups, and the GCSj was calculated for each pair of genomes, as described above.

To determine the optimal identity and mutual coverage cutoffs for clustering sequences, we evaluated the correlation between the GCSj calculated using the PanX orthology groups used before and the clusters and resulting P/A profiles obtained using different combinations of coverage and sequence identity cutoffs. We found that 80% identity with at least 70% bidirectional coverage was optimal or near-optimal across all four datasets, yielding the highest PCC in *P. aeruginosa*, *S. aureus*, and *H. pylori*, and a comparable value in *E. coli* (PCC 0.918 vs. 0.931 for the best combination; Fig. 3 and Supplementary Figs. S12–S13). GCSj values were overall robust to clustering thresholds across a broad identity/coverage range, supporting a single fixed cutoff across species.

**Figure 3.**
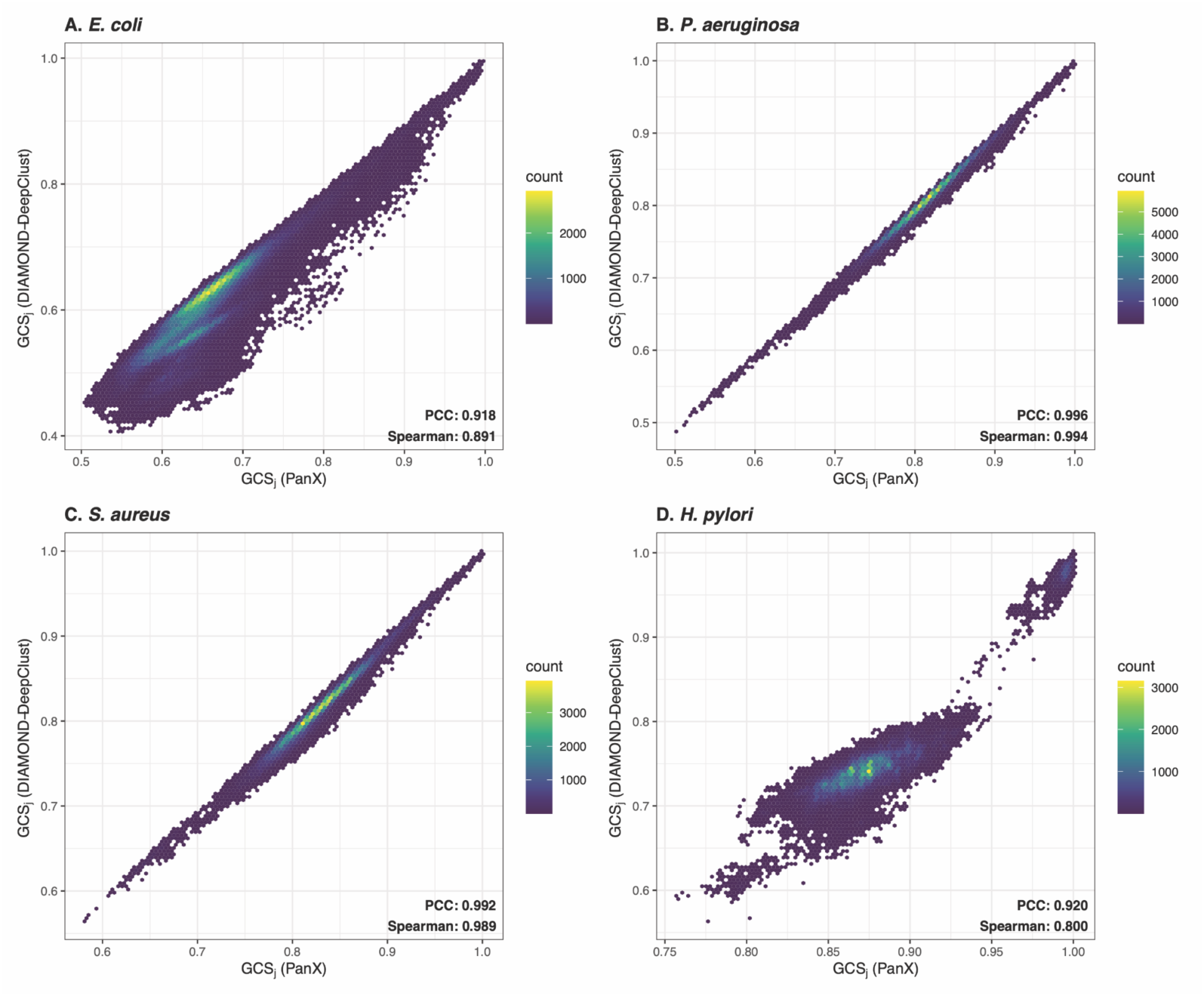
DIAMOND-DeepClust reproduces PanX-based GCSj across four bacterial species. GCSj calculated from PanX orthology groups strongly correlates with GCSj calculated from DIAMOND-DeepClust clusters (80% identity, ≥70% bidirectional coverage) in all four datasets (PCC ≥ 0.92), with the tightest agreement in *P. aeruginosa* and *S. aureus*. Each point represents a pairwise genome comparison, with color indicating point density (hexbin count). Pearson (PCC) and Spearman correlation coefficients between the two methods are shown for (A) *E. coli*, (B) *P. aeruginosa*, (C) *S. aureus*, and (D) *H. pylori*.

Based on these results, we suggest employing DIAMOND-DeepClust for fast and reliable estimation of gene-content similarity in bacteria. DIAMOND-DeepClust clusters proteins directly by sequence similarity, whereas tools such as PanX go a step further, refining these raw clusters into orthology groups using phylogenetic trees. This additional refinement dominates runtime: obtaining orthology groups with PanX required 1.9-11.2 hours per dataset (steps 03-06; 64-128 threads), with the tree-based refinement step alone accounting for 49–85% of the total running time. DIAMOND-DeepClust produced comparable clusters in under three minutes for every dataset at 64 threads, a 130–300× reduction in wall-clock time, while closely reproducing PanX-based GCSj values (PCC ≥ 0.92; Fig. 3, Supplementary Table S3).

### Clustering and Dereplicating Genomes by Gene Content Similarity

As PanGene-O-Meter proved to be highly effective in differentiating between genomes belonging to the same species, we tested whether it can also be used to dereplicate a set of bacterial genomes, so that one can nominate ‘representative’ genomes for downstream analyses and characterization. The naive approach is to calculate the gene-content similarity for all *n*(*n* − 1)/2 pairs of *n* genomes. The entries in such a similarity matrix can be clustered, and a representative from each cluster selected. This is a common practice for dereplicating a set of genomes using ANI (Olm et al. 2017).

However, computing and clustering all pairwise similarities is costly. Sketch-based pre-filtering can reduce this in principle. dRep, for example, first forms coarse primary clusters with Mash and computes exact ANI only within them (Olm et al. 2017), but within a single species most genomes fall into one primary cluster, so near-exhaustive comparison is still required. We thus propose GeneContRep, a greedy incremental clustering algorithm to dereplicate a list of genomes. GeneContRep adapts the greedy incremental clustering strategy of CD-HIT (Fu et al. 2012), originally developed for nucleotide and protein sequences, to gene presence/absence profiles, selecting representative genomes according to a user-defined gene-content similarity cutoff. Greedy representative selection has precedent in genome dereplication (e.g., dRep; Olm et al. 2017), though existing tools operate on nucleotide identity rather than gene content. Depending on the clusterability of the given data, this approach eliminates the need for comparisons of genomes already assigned early on to specific clusters. Only if no clusters are found for a given cutoff, all comparisons have to be calculated.

The GeneContRep algorithm operates as follows, given a set of genomes and a score cutoff:

1. Sort all genomes from longest to shortest (ties broken by genome identifier), giving a ranked list of unclustered genomes, *L*.
2. **Start a new cluster:** take the longest genome remaining in *L* as the cluster representative (*RepG*), and remove it from *L*.
3. **Populate the cluster:** for each remaining genome in *L*, compute its gene-content similarity (*GCSj* or *GCSo*) to *RepG*; if the score exceeds the cutoff, assign that genome to *RepG*’s cluster and remove it from *L*.
4. Repeat steps 2-3 until *L* is empty.

Sorting by length ensures the longest genome anchors each cluster, which is advantageous when using GCSo, since the overlap coefficient is normalized by the smaller genome and is therefore less penalized by genes missing from shorter assemblies (Eq. 2). GeneContRep supports both *GCSj* and *GCSo* as the clustering threshold and similarity measure.

We next compared GCSj-based clustering to ANI-based intra-species classification. A discontinuity in pairwise ANI has recently been reported at ∼99.2–99.8% ANI, with 99.5% proposed as a threshold delimiting intra-species units termed ‘genomovars’, which correspond closely to sequence types (Viver et al. 2024; Rodriguez-R et al. 2024; red band, Fig. 4). Figure 4 shows that clustering by gene content (GCSj) yields finer-grained clusters than ANI-based clustering, even at the high ANI cutoffs. This reflects the faster divergence of gene content relative to core-genome identity among closely related strains (Fig. 1B), rather than a difference between the clustering algorithms themselves. GCSj cutoffs of ∼0.80–0.90 yielded cluster numbers comparable, in most datasets, both to ANI-based clustering at this threshold and to the number of MLST sequence types. That is, gene content reached this level of resolution without requiring near-identical core genomes. This correspondence between GCSj and ST counts held even for *H. pylori*, where ANI-based clustering instead over-split the dataset markedly (>110 clusters for 21 STs already at 99% ANI), reflecting its high sequence diversity.

**Figure 4.**
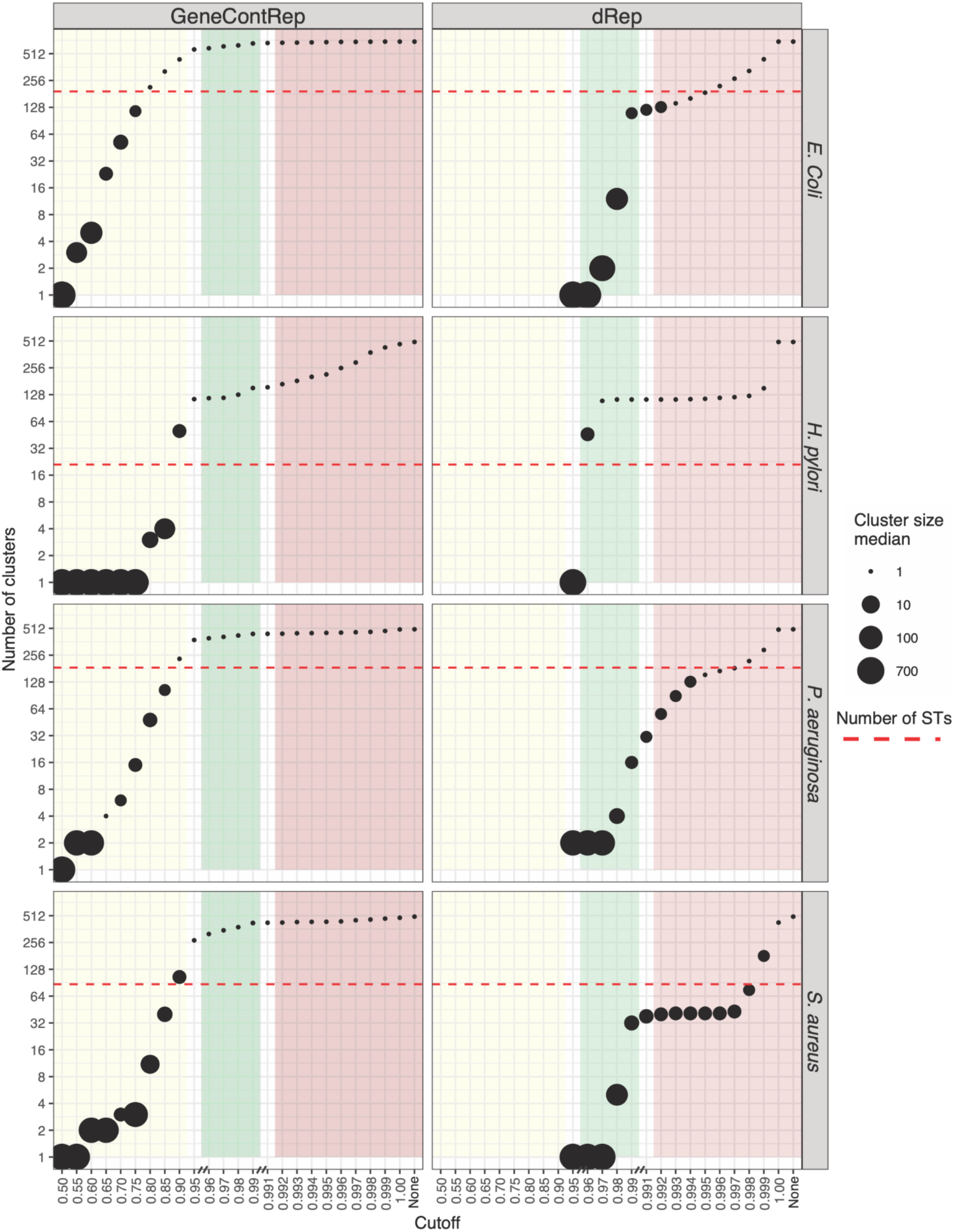
Gene-content-based clustering achieves higher resolution than ANI-based clustering, even within the ANI “genomovar” cutoff range. Number of clusters obtained across a range of similarity cutoffs using dRep (ANI-based clustering, right column) and GeneContRep (GCSj-based clustering, left column) for (A) *E. coli*, (B) *H. pylori*, (C) *P. aeruginosa*, and (D) *S. aureus*. Point size reflects the median number of genomes per cluster at each cutoff. The red dashed line indicates the number of sequence types (STs) predicted by the corresponding pubMLST scheme. The x-axis is broken to allow higher resolution within the densely sampled, near-identical genome range. Shaded regions indicate, from left to right: the cutoff range relevant for GCSj-based clustering (yellow), the “standard” ANI range of within-species variation (95-99%, green), and a range of high ANI cutoffs (99.0–99.9%, red) encompassing the proposed 99.5% genomovar threshold (Viver et al. 2024; Rodriguez-R et al. 2024).

We further applied this dereplicating approach to a set of *E. coli* genomes belonging to the B, E, and *Shigella* phylogroups (Supplementary Figure S14). The results demonstrate a clear gene-content-driven pattern grouping the different genomes. In all three cases, this grouping closely corresponds to the sub-phylogroup divisions suggested by MASH analysis (Abram et al. 2021).

### Gene-Content Similarity Captures Clinically Relevant Functional Diversity

Having established that GCSj provides finer resolution than ANI for genome classification and dereplication, we next asked whether this resolution is useful for capturing meaningful biological and clinical variation. We analyzed the relationship between genome similarity and functional diversity across all four species datasets. Using DRAM-derived KEGG functional profiles, we calculated pairwise accessory functional distances (Jaccard) and compared them against ANI and GCSj (Supplementary Figure S15). Even among highly similar genomes (ANI > 99.5%), ANI-defined groups retained substantial functional variation, with elevated median accessory functional distances across all ANI bins. In contrast, functional distance decreased sharply and consistently with increasing GCSj across all four species, with the highest GCSj bin (>0.95) showing markedly lower functional distances than any ANI bin tested. This pattern was consistent across species with very different pan-genome architectures, from the open pan-genome of *E. coli* to the relatively closed pan-genome of *H. pylori* (though some H. pylori ANI bins are sparsely populated; Fig. S15), indicating that GCSj captures a dimension of functional organization that ANI alone cannot resolve.

To test whether gene-content-based dereplication preserves clinically relevant diversity better than standard ANI-based approaches, we used experimentally determined antimicrobial resistance phenotypes as an external ground truth, determined independently of any genome-similarity metric. We constructed a dedicated validation dataset of 1,000 high-quality *Klebsiella pneumoniae* genomes from two major multidrug-resistant lineages, ST258 and ST307 (500 genomes each), sourced from the BV-BRC (Bacterial and Viral Bioinformatics Resource Center) database. These genomes were selected based on high assembly quality and the availability of matched experimental antimicrobial resistance (AMR) phenotypes. Genomes were sampled to ensure balanced representation of carbapenem-resistant and carbapenem-susceptible isolates within each ST, providing a stringent test of whether similarity metrics can discriminate phenotypically relevant genomic variation.

We first assessed whether GCSj could resolve phenotypic divergence within the narrow range of near-identical genomes (ANI ≥ 99.9%) where nucleotide identity alone provides limited resolution. Restricting our analysis to genome pairs within this range, AMR profile divergence decreased significantly with increasing GCSj threshold, with the effect size strengthening monotonically and reaching a moderate rank-biserial correlation (r = 0.38) at GCSj ≥ 98% in both lineages (Fig. 5A). The clinical relevance of this discrimination was further illustrated by examining carbapenem resistance concordance. The percentage of genome pairs phenotypically discordant for carbapenem resistance dropped substantially under progressively stricter GCSj thresholds, falling from 45% at the ANI ≥ 99.9% baseline to just 13% at GCSj ≥ 98% in ST258, and from 55% to 30% in ST307 (Fig. 5B). These results indicate that GCSj captures resistance-relevant gene-content variation that remains invisible to nucleotide-based comparisons.

**Figure 5.**
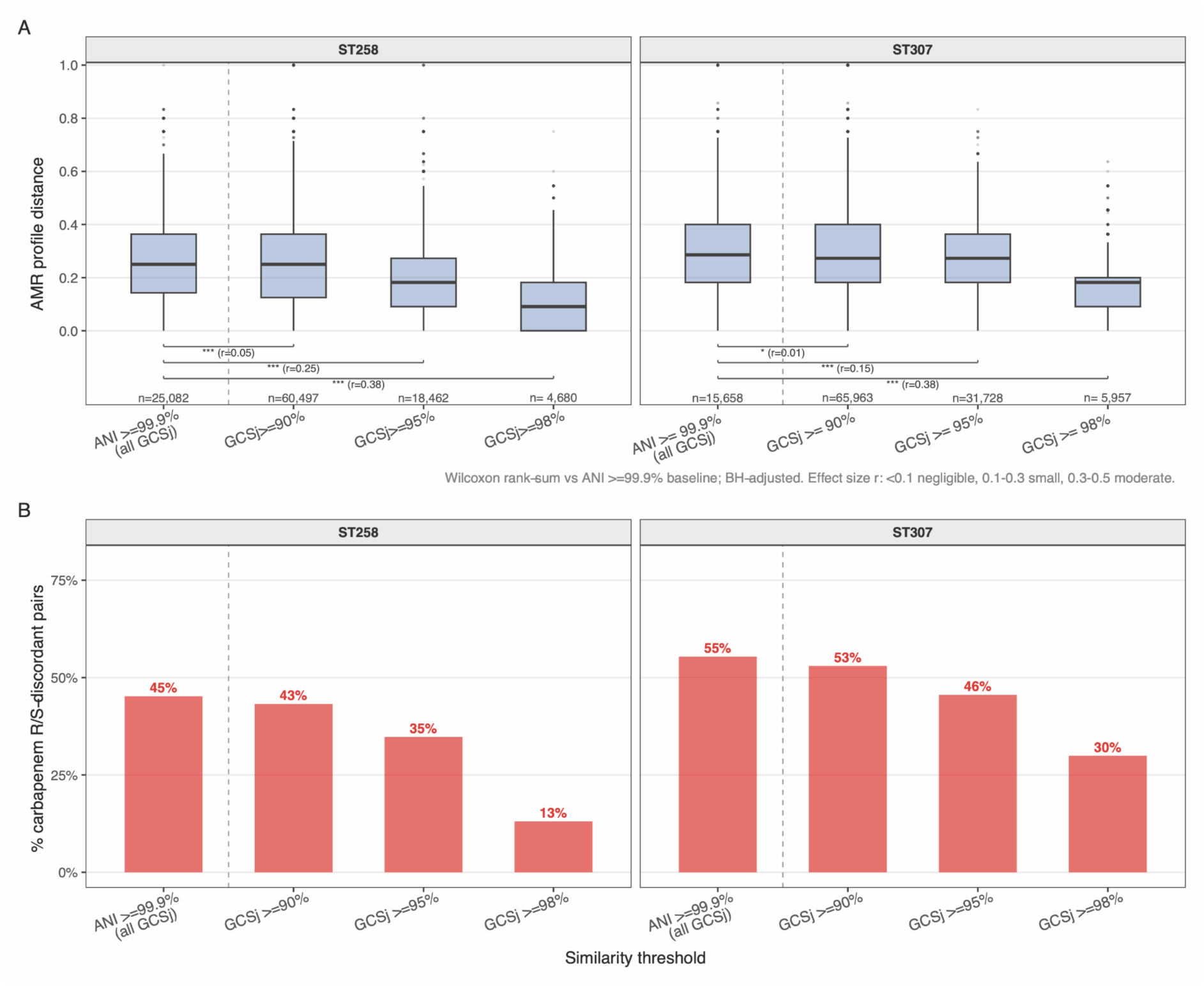
GCSj resolves AMR-relevant variation within highly similar *Klebsiella pneumoniae* genome pairs. Analyses restricted to genome pairs with ANI ≥ 99.9% are shown separately for lineages ST258 and ST307. The dashed vertical line separates the ANI-only baseline (all GCSj values) from progressively stricter GCSj filters. (A) Distribution of pairwise AMR profile distances (Jaccard) across similarity thresholds. Boxes show median and IQR; whiskers extend to 1.5×IQR; outliers shown as individual points. Numbers above each box indicate the number of genome pairs. Brackets indicate Wilcoxon rank-sum tests against the ANI ≥ 99.9% baseline (BH-adjusted; *** p < 0.001, * p < 0.05); rank-biserial correlation r is shown as effect size (r < 0.1 negligible, 0.1-0.3 small, 0.3-0.5 moderate). Given the large number of pairwise comparisons, statistical significance is expected even for negligible effects; we therefore emphasize effect sizes (r) over p-values. (B) Percentage of genome pairs that are phenotypically discordant for carbapenem resistance (one isolate resistant, one susceptible) across the same similarity thresholds. In ST258, discordant pairs decrease from 45% at the ANI baseline to 13% at GCSj ≥ 98%, demonstrating that stricter gene-content thresholds substantially enrich phenotypically concordant genome pairs.

We next asked whether the k-mer-based accessory distance estimated by PopPUNK (Lees et al. 2019), a widely used tool for bacterial epidemiology, could provide equivalent resolution within these lineages. Between-ST comparisons were clearly separated from within-ST comparisons by core distance (core distance ≈ 0.008 between STs versus ≈ 0 within STs), confirming that PopPUNK correctly resolves the distinct evolutionary trajectories of the two lineages. However, within-ST accessory distances spanned a broad, continuous range (approximately 0-0.15, with a diffuse tail extending to ∼0.2) with no clear discrete substructure in either ST258 or ST307, demonstrating that PopPUNK operates effectively at between-lineage resolution although it cannot discriminate the within-ST gene-content variation relevant to AMR phenotype prediction (Supplementary Figure S16). This is consistent with PopPUNK’s design as a population-level epidemiological classifier rather than providing a fine-grained gene-content similarity metric. In contrast, GCSj, by relying on explicit orthology group assignments, resolves within-ST diversity at the resolution needed to recover clinically relevant variation.

Finally, we applied GeneContRep to select representative genomes from the *K. pneumoniae* dataset at GCSj thresholds of 90%, 95%, and 98%, and compared the retained diversity against standard dRep dereplication at 99.9% ANI (Fig. 6A). ANI-based dereplication at 99.9% reduced each ST to 17 (ST258) and 11 (ST307) representative genomes, but at severe cost to diversity: these representatives captured only a small fraction of the capsule (KL) types, O-locus (OL) types, and carbapenemase variants present in the full cohort. Gene-content-based dereplication retained substantially more clinically relevant diversity at all tested thresholds. In ST307, GCSj at 95% selected 60 representative genomes, approximately five times more than ANI 99.9%, while capturing 100% of the KL types, 90% of the OL types, and a substantially larger proportion of carbapenemase variants including both *bla*KPC and *bla*NDM alleles. A comparable pattern was observed in ST258, where GCSj at 95% selected 73 representatives and recovered diversity largely absent from the 17 ANI-selected representatives (Fig. 6A). We note that we used 99.9% ANI as the dRep threshold because this is the highest value considered reliable by the tool’s authors; above ∼99.9% ANI, values approach the limit of detection, since even self-comparisons typically yield ∼99.99% ANI (Olm et al. 2017; dRep Documentation). ANI-based dereplication therefore cannot be pushed to finer resolution, whereas gene content continues to resolve structure well beyond this point.

**Figure 6.**
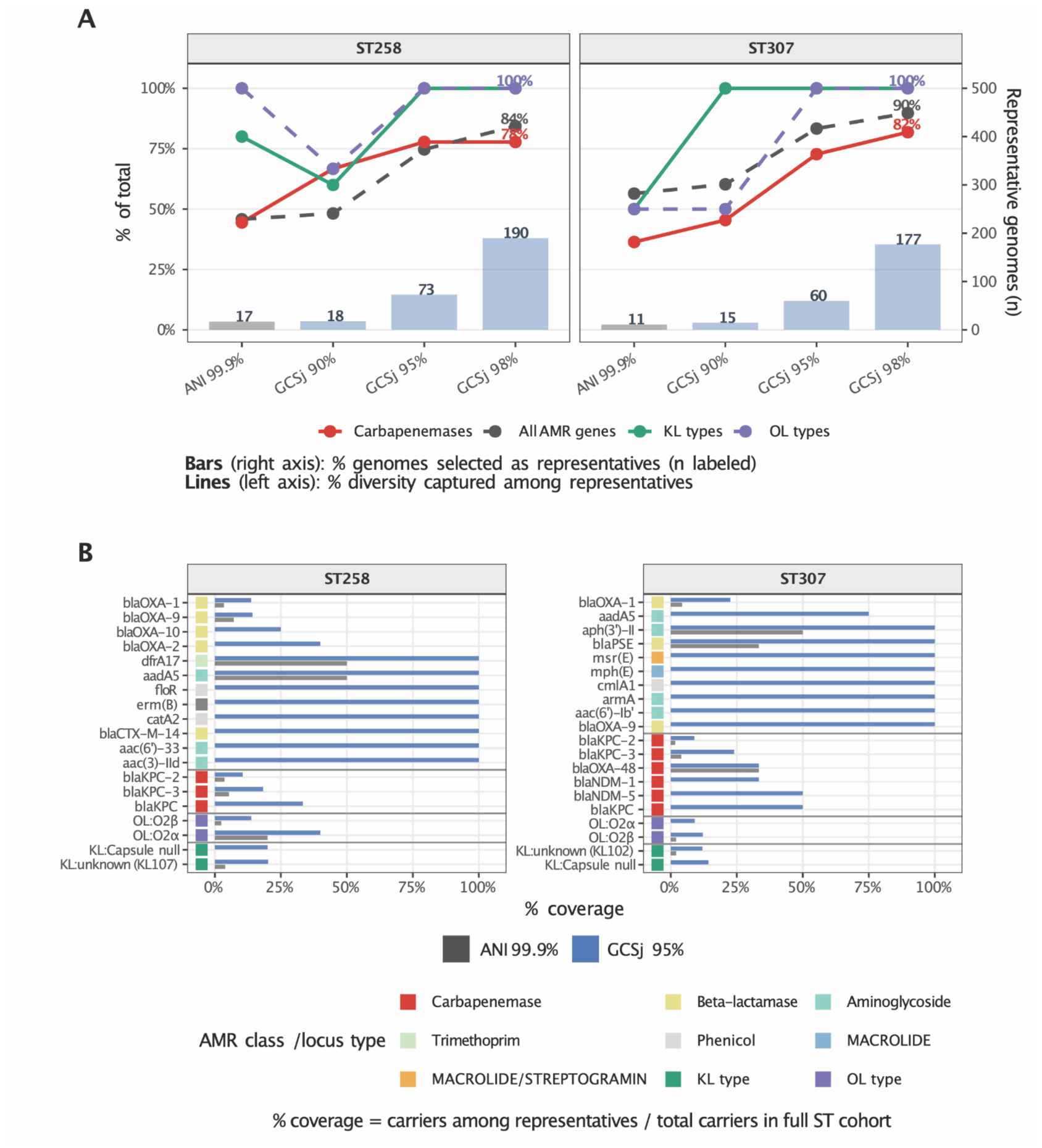
GCSj-based dereplication retains substantially more clinically relevant diversity than ANI-based dereplication in *Klebsiella pneumoniae*. Analyses are shown separately for lineages ST258 and ST307 (500 genomes each). (**A**) Bars (right axis) show the number of representative genomes selected by ANI-based dereplication (99.9%) and GCSj-based dereplication (90%, 95%, and 98%). Lines (left axis) show the fraction of total diversity captured among representatives for four feature classes: carbapenemase determinants (red), all AMR determinants (black dashed), capsule KL types (green), and O-locus OL types (purple dashed). In ST307, GCSj at 95% selects 60 representatives and captures 100% of KL types, 90% of OL types, and 82% of carbapenemase variants, compared to just 11 representatives under ANI 99.9%. (**B**) Per-feature coverage reveals specific determinants missed by ANI and preserved by GCSj. Bars show the percentage of genomes carrying each feature that are represented among ANI 99.9% representatives (grey) or GCSj 95% representatives (blue), relative to all carriers in the full ST cohort. Features shown include all carbapenemases, the most differentially covered non-carbapenem AMR determinants, and the most differentially covered KL and OL locus types. Colored squares indicate AMR class or locus type.

A curated feature-level analysis makes explicit what ANI misses and GCSj preserves (Fig. 6B; Supplementary Figure S17). Among the ANI 99.9% representatives, coverage of carbapenemase genes — including *blaKPC-2*, *blaKPC-3*, *blaNDM-1*, and *blaOXA* variants — was near zero or severely incomplete in both STs. GCSj 95% representatives captured these clinically relevant determinants at substantially higher coverage, often exceeding 75-100% of carriers in the full cohort. Beyond carbapenemases, GCSj representatives also retained a broader spectrum of beta-lactamase, aminoglycoside, and other resistance determinants, as well as a more complete representation of KL and OL locus diversity - features directly relevant to vaccine design, serotyping, and infection management. The comprehensive per-feature comparison in Supplementary Figure S17 further confirms that this improvement is consistent across both STs and across all major AMR classes represented in the dataset.

Altogether, these results establish that gene-content similarity, as implemented in PanGene-O-Meter, captures biologically and clinically meaningful dimensions of intra-species diversity that remain hidden when genomes are compared by nucleotide identity alone. GeneContRep provides an effective and tuneable framework for reducing dataset size while faithfully preserving the functional, phenotypic, and clinically relevant diversity of the original population.

## DISCUSSION

Dereplicating genomes and selecting a subset of ‘representative genomes’ is a common practice in comparative genomics analyses (Perrin and Rocha 2021; Shalon et al. 2023; Olm et al. 2017). Here we suggest an alternative that directly groups genomes by their gene-content. Grouping genomes by gene-content should benefit comparative genomics studies by allowing us to focus on intra-species gene-content diversity, thereby enhancing our understanding of the role of mobile or variable genetic elements in bacterial evolution. We have already demonstrated the usefulness of an earlier version of this approach for functional exploration of diversity among *Pseudomonas* spp. isolates from wild *Arabidopsis thaliana* plants (Shalev et al. 2022).

The foundation of PanGene-O-Meter lies in how the pangenome is constructed. Major errors in orthology-group assignment can result in both overestimation and underestimation of GCS. Theoretically, assigning diverged genes erroneously to the same orthology group will result in higher (over-estimated) GCS, as it implies a greater number of common genes. Conversely, excessive splitting of orthology groups will result in lower (under-estimated) GCS, as it implies fewer common genes. We demonstrated that gene-clusters provided by DIAMOND-DeepClust (Buchfink et al. 2026) using ‘one-fits-all’ clustering parameters can still provide an accurate estimate of GCS similar to the more laborious and careful assignment of genes into orthology groups as done by PanX (Ding et al. 2018). This suggests that the effect of errors in orthology-group assignment is overall minor, and the overall pattern is robust for GCS estimation. We implemented in PanGene-O-Meter an advanced option to exclude ‘private’ orthology-groups (present in a single taxon), which are the most likely to arise from assembly errors or under-clustering. In our datasets, enabling this option changed GCS estimates only marginally, indicating that private groups contribute little to the overall signal. Another approach to assess the robustness of the GCS estimation is to use different orthology-group assignment methods. We note that a possible bias for GCS estimation could be introduced by partial or highly contaminated genomes. We provide both GCSj and GCSo metrics, which can help to reduce the impact of such biases. Additionally, it is advisable to estimate the quality of the analyzed genomes using metrics such as CheckM2 (Chklovski et al. 2023) and BUSCO (Tegenfeldt et al. 2025) and exclude low-quality genomes from the analyses (or at least to treat them with caution).

Gene-content similarity was suggested before as a means to classify viruses (Roux et al. 2021) and to study their evolution (Yutin et al. 2013). In the context of bacterial genomes it has been mostly used for comparing ‘functional profiles’ based on a set of predefined functions (provided, for example, by COG or KEGG) (Hernández-González et al. 2018) or based on the assignment of genes to sets of predefined orthology groups (e.g., by eggNOG) (Tu and Lin 2016; Maistrenko et al. 2020). Here we generalize this concept and provide a comprehensive and efficient pipeline for quantifying gene-content similarity as well as dereplicating a set of bacterial genomes based on their gene-content. This is especially useful for characterizing intra-species diversity.

A natural question is how PanGene-O-Meter compares to other tools that estimate accessory genome distances. The most relevant is PopPUNK (Lees et al. 2019), which uses k-mer distributions to jointly estimate core and accessory distances for population-scale epidemiological classification. PopPUNK is highly effective at between-lineage resolution and has become a standard tool for bacterial outbreak tracking and lineage assignment. Comparing PopPUNK with gene content similarity measures of *Klebsiella pneumoniae* ST258 and ST307 revealed an important distinction in resolution scale: within highly clonal lineages, where core distances approach zero, PopPUNK accessory distances span a continuous range with no resolvable substructure, effectively collapsing the entire within-ST diversity into a single undifferentiated cluster. In contrast, GCSj, by relying on explicit orthology group assignments rather than k-mer distributions, retains the power to discriminate within-ST gene-content variation, which is directly relevant to carbapenem resistance phenotype and capsule locus diversity. This difference reflects a fundamental difference in purpose and design philosophy: PopPUNK is optimized for population-level lineage assignment across a broad range of genomic diversity, whereas PanGene-O-Meter is designed specifically for fine-grained intra-lineage comparisons, where accessory genome variation is the primary signal of interest. The two approaches are therefore complementary rather than competing: PopPUNK excels at assigning isolates to lineages and tracking transmission at the epidemiological scale, while PanGene-O-Meter provides the resolution needed to characterize functional and phenotypic diversity within those lineages. We suggest that a natural workflow would use PopPUNK for initial lineage assignment followed by PanGene-O-Meter for within-lineage functional diversity analysis and representative genome selection, combining the computational efficiency of k-mer approaches with the biological interpretability of explicit gene-content comparison.

With respect to threshold selection, GCSj cutoffs between ∼0.80 and 0.90 gave resolution broadly comparable to sequence-type boundaries and the ANI genomovar zone across the species tested here. This makes GCSj ≈ 0.90 a practical starting point for dereplication in the absence of prior knowledge, as it yielded at least as many clusters as sequence types in all four datasets. For datasets containing incomplete assemblies or metagenome-assembled genomes, we recommend GCSo over GCSj, which in our simulations remained robust to up to 30% gene-content incompleteness, particularly when comparing a complete genome to an incomplete one. Users seeking finer resolution, for example, to discriminate phenotypically distinct strains within a single sequence type, as shown here for carbapenem-resistant *K. pneumoniae*, should consider stricter GCSj thresholds of 0.95-0.98, where gene-content differences become increasingly predictive of functional and phenotypic divergence. Because resistance phenotype is measured independently of genome sequence similarity, this provides external validation that GCSj resolves biologically meaningful variation, rather than being assessed only against gene content itself. We note that our clinical validation focused on carbapenem resistance in two *K. pneumoniae* lineages; extending this analysis to other pathogens, resistance phenotypes, and clinically relevant traits will be important to establish the generality of gene-content-based dereplication for surveillance applications.

The potential application of methods such as PanGene-O-Meter is not limited to bacterial genomes. For example, recent studies have demonstrated that fungal gene-content can be highly variable (Perrier and Barber 2024). Notably, such variation has also been observed among closely related species (e.g., (Sato et al. 2025)). A major requirement for the application of PanGene-O-Meter to non-bacterial genomes is the availability of accurate and efficient gene-annotation and orthology-group assignment methods. The development of advanced deep-learning algorithms for eukaryotic gene annotations, such as Helixer (Holst et al. 2023) and ANNEVO (Ye et al. 2025), and advanced orthology-group algorithms, such as OrthoFinder3 (Emms et al. 2026), is expected to significantly enhance the accuracy and efficiency of these tasks. By combining these methods and leveraging the growing availability of fully assembled eukaryotic genomes, we anticipate that systematic studies of gene-content diversity and its functional implications in eukaryotes will become feasible in the near future. This would represent a direct extension of PanGene-O-Meter to provide insights into the evolution and diversity of eukaryotic genomes.

In conclusion, we present PanGene-O-Meter as a framework for quantifying gene-content similarity among bacterial genomes. In addition, we have implemented an efficient clustering algorithm, GeneContRep, which clusters and dereplicates genomes based on their gene-content. This dereplication step yields a more compact set of representative genomes, enabling more focused downstream functional analyses. By providing a direct estimate of gene-content variability, PanGene-O-Meter can facilitate, for example, the identification of genes that differentiate bacteria sampled from different environments, thereby enhancing comparative-genomics studies.

## Methods

### Data Collection and Preprocessing

Genome assemblies of *H. pylori* were downloaded from the AllTheBacteria database (Hunt et al. 2024). Gene calling was performed using the PROKKA pipeline [version 1.14.6] (Seemann 2014) with default parameters. Genome assemblies and gene annotations of *E. coli*, *S. aureus,* and *P. aeruginosa* were obtained from NCBI-Genomes (Sayers et al. 2025). Assemblies for all four species were evaluated with CheckM2 (Chklovski et al. 2023), and only those with completeness over 95% and contamination less than 5% were retained. The full list of accession IDs is provided in Supplementary Table S2.

### Sequence Typing and Genome Similarity

Sequence types (STs) were assigned using the mlst program [version 2.23] (Seemann 2022) and the PubMLST database from August 2024 (Jolley and Maiden 2010). Genome similarity was assessed using the following methods:

- **ANI** was calculated using skani v0.2.2 with default parameters (Shaw and Yu 2023). Genome dereplication was performed with dRep v3.5.0, which clusters genomes using its internal fastANI implementation (Olm et al. 2017); skani was used for all other ANI computations reported here.
- **SynTracker** scores were calculated using SynTracker [version 1.3] (Enav et al. 2024) with reference genomes from GTDB (accessions GCF_003697165.2, GCF_001457615.1, GCF_900478295.1, and GCF_001027105.1). For *E. coli* and *P. aeruginosa*, 1.2 Mb (∼20%) of the GTDB reference genomes was used to speed calculation. For all datasets, SynTracker run with the 95% identity cutoff was used. The average synteny score across 100 regions was used as the SynTracker score for each pair of genomes.
- **PopPUNK** Pairwise core and accessory genomic distances were estimated using PopPUNK v2.7.8 (Population Partitioning Using Nucleotide K-mers) (Lees et al. 2019). A sketch database was constructed using the *--create-db* function with default k-mer settings, a maximum accessory distance threshold of 1.0 (*--max-a-dist 1.0*). The resulting pairwise distance matrices, stored in binary format (*.dists.pkl* and *.dists.npy*), were parsed using a custom Python script. Sample identifiers were retrieved from the pickle file, and the corresponding core and accessory distance values were extracted from the NumPy array. PopPUNK decomposes genomic distances into two components: core distance (divergence in conserved regions) and accessory distance (differences in flexible genome content). We used the accessory-distance-derived similarity for comparisons with GCSj; core distance is shown only to confirm lineage separation in the *K. pneumoniae* analysis (Fig. S16).
- **The percentage of unaligned base-pairs** for each pair of genomes was calculated using DNAdiff version 1.3 (Kurtz et al. 2004).
- **Mobile elements distance.** Mobile elements were annotated using geNomad [version 1.5.2] (Camargo et al. 2024), AMRFinderPlus [version 3.10.16] (Feldgarden et al. 2021), and IslandPath-DIMOB [version 1.0.6] (Bertelli and Brinkman 2018). Next, for each dataset, all mobile elements were clustered using vConTACT2 [version 0.11.3] (Bin Jang et al. 2019). Finally, the Jaccard similarity was calculated for all genome pairs utilizing their mobile elements presence-absence profiles.

### Simulation of Genome Incompleteness

To assess the sensitivity of our similarity measures to assembly incompleteness, we simulated random gene absences using the *E. coli* phyletic-pattern matrix. For 20,000 randomly selected genome pairs, we artificially degraded the assemblies by randomly changing a subset of “present” genes (1) to “absent” (0). We tested two scenarios: (i) degrading both genomes in the pair, and (ii) degrading only one genome in the pair. Incompleteness was simulated at levels of 1%, 5%, 10%, 20%, and 30% of total gene count, with 10 independent replicate draws per pair at each level. For each degraded pair, GCSj and GCSo were recalculated, and the absolute error was determined by comparing the resulting values against the ground-truth similarity computed from the original, complete genomes.

### Nested ANI-GCSj similarity heatmaps

For visualization of the relationship between ANI and GCSj similarity, pairwise ANI and GCSj matrices were ordered using a nested clustering strategy. First, genomes were clustered by hierarchical clustering of ANI-derived distances, defined as 100-ANI, using average linkage. The resulting ANI dendrogram was cut into ten coarse groups for visualization. Within each ANI-defined group, genomes were reordered by hierarchical clustering of GCSj-derived distances, defined as 100−GCSj. This final genome order was applied to both the ANI and GCSj matrices, allowing direct comparison of nucleotide similarity and gene-content similarity in the same genome order. The fixed number of ANI groups was used only for visualization and was not interpreted as a biological cutoff.

### Phylogenetic trees

Maximum likelihood trees and distances were inferred using IQ-TREE [version 3.0.1] (Wong et al. 2025). Gene-content trees were inferred from the phyletic pattern using the best-fit BIN model (-st BIN), selected automatically by ModelFinder (Kalyaanamoorthy et al. 2017). Core-genome trees were inferred from a concatenated nucleotide alignment of orthology groups present in a single copy in 100% of the analyzed genomes (strict core), as produced by PanX (Ding et al. 2018) and aligned with MAFFT (Katoh and Standley 2013), using the GTR substitution model with empirical base frequencies and FreeRate rate heterogeneity (-m GTR+F+R) (Yang 1995; Soubrier et al. 2012). The strict core comprised 241 genes (153,036 aligned positions) for *E. coli*, 914 genes (630,972 positions) for *P. aeruginosa*, 775 genes (799,050 positions) for *S. aureus*, and 541 genes (517,596 positions) for *H. pylori*. The smaller core of *E. coli* reflects the deliberate inclusion of all phylogroups, which maximizes intra-species diversity in that dataset. Trees and presence–absence matrices were plotted using the ggtree package in R (Yu 2020).

### Pan-genome Construction

The pan-genome was constructed using PanX version 1.5.1 (Ding et al. 2018) with the following parameters: -dmdc -dcs 50 -sitr. DIAMOND version 2.1.9 (Buchfink et al. 2026) was used with the following parameters: cluster --approx-id *ID* --mutual-cover *COVER* -- masking 0 --soft-masking 0 --comp-based-stats 0 -M 64G; where *ID* and *COVER* are the clustering percent identity and mutual coverage cutoffs.

### DRAM-derived KEGG functional distance

To evaluate whether the granularity provided by gene-content similarity reflects meaningful functional biological variation, we analyzed the functional repertoires of the four primary datasets (*E. coli*, *H. pylori*, *P. aeruginosa*, and *S. aureus*). Protein sequences derived from the assemblies were annotated using DRAM (Distilled and Refined Annotation of Metabolism) v1.5.0 with default parameters (Shaffer et al. 2020). KEGG Orthology (KO) assignments were extracted from the distilled DRAM annotations to construct a binary presence/absence functional profile for each genome. To ensure the distance metric specifically captured functional plasticity rather than baseline taxonomic similarity or transient sequencing noise, we restricted our analysis to the accessory functional genome. Specifically, KO terms present in greater than 95% of genomes (core functions) or fewer than 5% of genomes (rare/transient functions) within each species dataset were excluded. Pairwise accessory functional divergence was then quantified by calculating the Jaccard distance between the filtered accessory KO profiles for all genome pairs.

### Klebsiella pneumoniae Clinical Validation

A cohort of 1,000 multidrug-resistant *Klebsiella pneumoniae* genomes (500 each of ST258 and ST307) was obtained from the BV-BRC database (Shukla et al. 2026). Assembly quality was evaluated using CheckM2 v1.1.0 (Chklovski et al. 2023), and genomes were annotated with PROKKA v1.14.6 (Seemann 2014). Capsule (KL) and O-antigen (OL) locus types were assigned using Kleborate v3.2.4 (Lam et al. 2021). Antimicrobial resistance (AMR) genes were annotated with AMRFinderPlus v3.10.16 (database version 2024-07-22.1). Pairwise AMR profile divergence was calculated as the Jaccard distance derived from the binary presence/absence matrix of all identified AMR determinants. Pairwise genomic distances within each ST were computed using skani v0.2.2 (for ANI), PanGene-O-Meter relying on PanX (for GCSj), and PopPUNK v2.7.8 with default k-mer distributions (for core/accessory distances). Sequence-based genome dereplication was performed using dRep (fastANI algorithm) at a strict primary ANI threshold of 99.9%, utilizing CheckM2 metrics to score representative genomes. Gene-content-based dereplication was executed using GeneContRep at GCSj thresholds of 90%, 95%, and 98%. Experimental antimicrobial susceptibility phenotypes (resistant/susceptible calls) were obtained from the BV-BRC metadata accompanying each genome. A genome pair was considered phenotypically discordant for carbapenem resistance when one isolate was recorded as carbapenem-resistant and the other as carbapenem-susceptible.

### GeneContRep implementation

GeneContRep is implemented as part of the PanGene-O-Meter package (https://github.com/HaimAshk/PanGene-O-Meter); it performs greedy incremental clustering as described in Results, using GCSj or GCSo as the similarity measure.

### Statistical analysis

All statistical analyses were performed in R (v4.4.0). Correlations between similarity measures are reported as Pearson (PCC) and Spearman coefficients. Heteroscedasticity of GCSj across ANI bins was tested with the Fligner–Killeen test. Differences in AMR profile distance across similarity thresholds were assessed with two-sided Wilcoxon rank-sum tests against the ANI ≥ 99.9% baseline, with Benjamini-Hochberg correction for multiple comparisons and rank-biserial correlation (r) as effect size, all computed using the rstatix package (v1.0.0). Effect sizes are reported as the rank-biserial correlation (r), interpreted as negligible (<0.1), small (0.1-0.3), or moderate (0.3-0.5). Given the large number of pairwise comparisons, statistical significance is expected even for negligible effect sizes; we therefore emphasize effect sizes over p-values throughout.

## Data and Code Availability

PanGene-O-Meter is available at: https://github.com/HaimAshk/PanGene-O-Meter. The constructed pan-genome for each species and all calculated distances are available at ZENODO repository: doi:10.5281/zenodo.17025330.

## Author Contributions

HA designed and executed the project. DW supervised the project. HA prepared the first draft of the manuscript. HA and DW finalized the manuscript.

## Competing Interests

DW holds equity in Computomics, which advises plant breeders. DW in the past consulted for KWS SE, a globally active plant breeder and seed producer. HA declares no competing interests.

## Funding

HA was supported by the Alexander von Humboldt Foundation. This study was supported by the Max Planck Society and the Novozymes Prize of the Novo Nordisk Foundation (DW).

## Acknowledgments

We thank Hajk-Georg Drost and Benjamin Buchfink for guidance in the use of DIAMOND DeepClust. We thank Sheila Roitman, Gal Ofir, Miriam Lucke, Hagay Enav, David Burstein, and Tal Pupko for feedback on the manuscript.

## Notes

### Summary of Updates

This major revision introduces extensive new analyses to demonstrate the immediate clinical, functional, and ecological utility of PanGene-O-Meter. Key additions include: 1. Clinical Validation (Klebsiella pneumoniae): We analyzed a curated cohort of 1,000 multidrug-resistant genomes (ST258 and ST307). We demonstrate that standard ANI-based deduplication (at 99.9%) actively erases clinically vital diversity, including carbapenemases and capsule (KL)/O-locus types. In contrast, our GCSj metric preserves this diversity and accurately predicts phenotypic AMR divergence. 2. Functional Pangenomics (DRAM/KEGG): We added comprehensive functional profiling, proving that gene-content similarity captures accessory metabolic and functional plasticity that core-sequence identity (ANI) obscures. 3. Robustness to MAGs: We introduced extensive gene-loss simulations demonstrating that our overlap-based metric (GCSo) remains uniquely robust against genome fragmentation, making it highly suitable for comparing reference genomes to incomplete metagenome-assembled genomes (MAGs). 4. Enhanced Benchmarking: We included direct comparisons to network-based sub-lineage clustering (PopPUNK), further validating the superior sub-lineage resolution provided by PanGene-O-Meter.

