## Supplementary information for "PanGene-O-Meter: Intra-Species Diversity Based on Gene-Content"

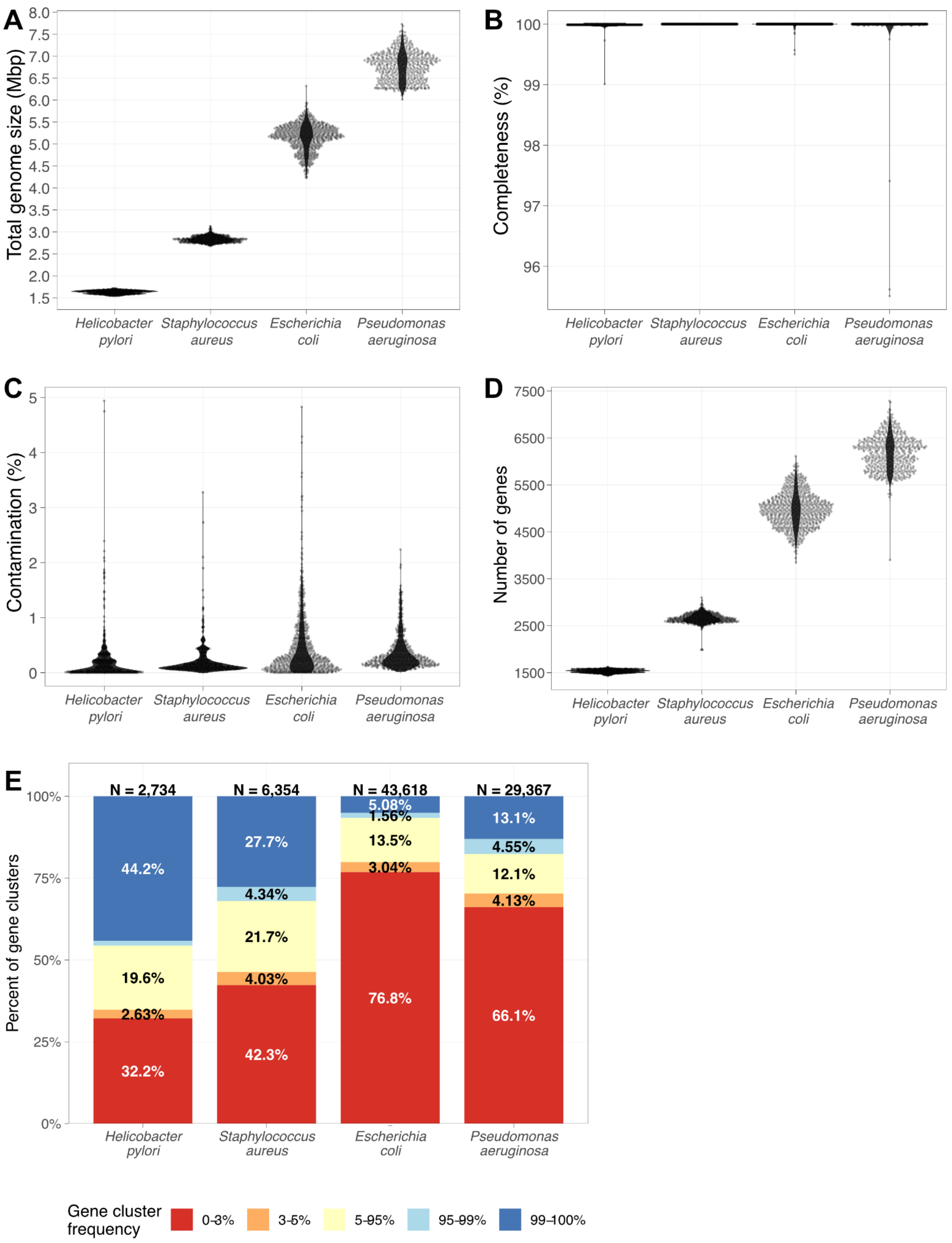


**Figure S1. Quality of the analyzed genomes and intra-species gene-content variability.** Distribution of (A) Total genome size; (B) Completeness; (C) Contamination; (D) The number of protein-coding genes per species. (E) Pan-genome view of the gene-content variability and orthology groups frequency. Most of the genes are part of the accessory genome (present in fewer than 95% of strains).

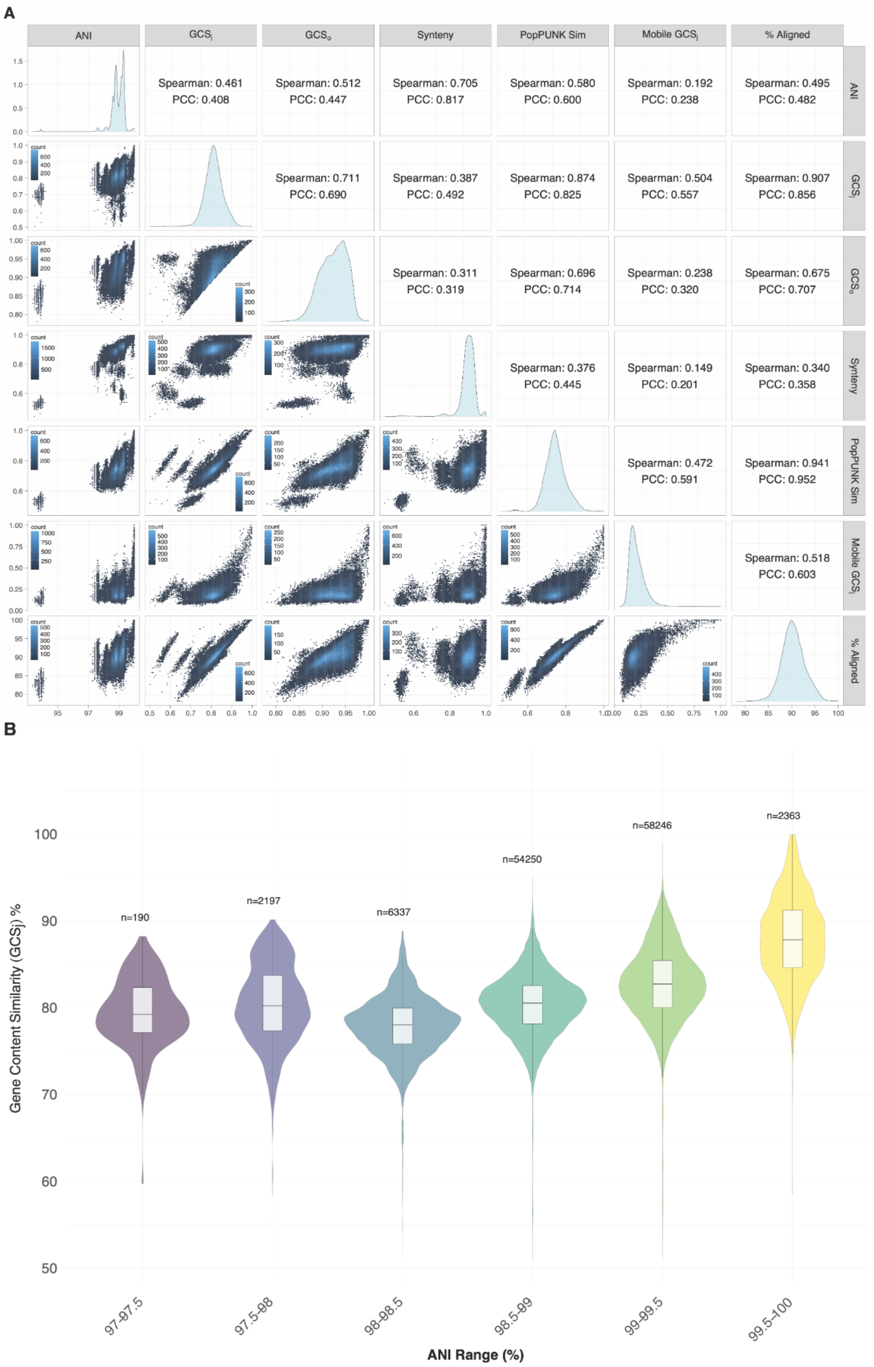


**Figure S2 - Comparing pairwise similarity measures for the *Pseudomonas aeruginosa* datasets. (A)** Pairwise comparisons of similarity measures: ANI; gene-content similarity by Jaccard (GCSj) and overlap coefficient (GCSo); synteny score (SynTracker); PopPUNK accessory similarity; mobile-element profile similarity (Mobile GCSj); and fraction of aligned bases (% Aligned). Upper-right panels give Pearson (PCC) and Spearman coefficients. Gene-content similarity varies widely even among pairs with near-identical ANI, as quantified in (B). **(B)** Distribution of GCSj values across binned ANI ranges. Violin plots show the density distribution of GCSj, with internal boxplots indicating the median and interquartile range (IQR). The number of pairwise comparisons (*n*) is shown above each bin. The variance of GCSj is not homogeneous across ANI ranges (Fligner-Killeen test, p-value < 2.2e-16), and the spread of gene-content similarity peaks at the highest sequence identities (ANI 99.5-100%), highlighting extensive accessory genome variation among closely related strains.

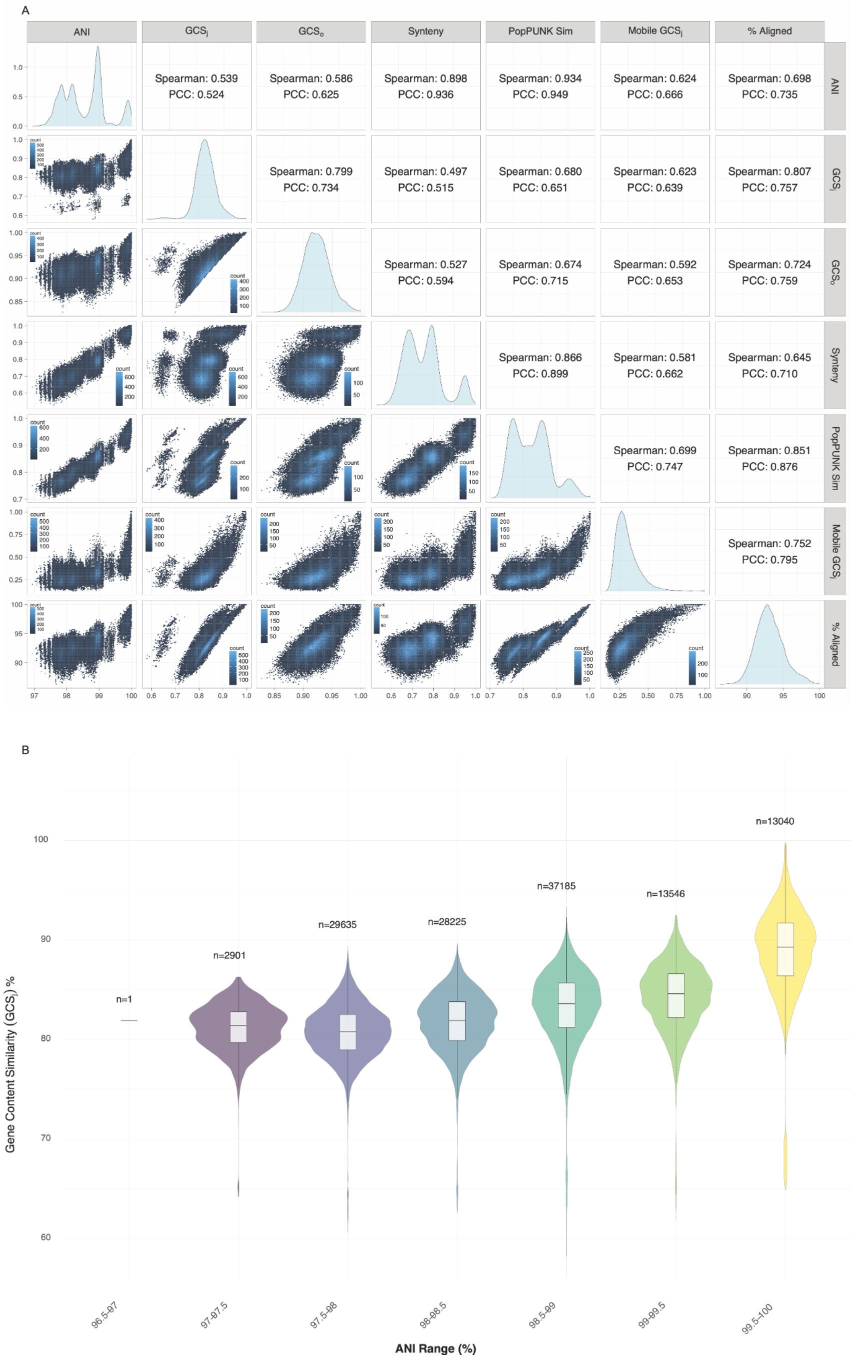


**Figure S3 - Comparing pairwise similarity measures for the *Staphylococcus aureus* datasets.** **(A)** Pairwise comparisons of similarity measures: ANI; gene-content similarity by Jaccard (GCSj) and overlap coefficient (GCSo); synteny score (SynTracker); PopPUNK accessory similarity; mobile-element profile similarity (Mobile GCSj); and fraction of aligned bases (% Aligned). Upper-right panels give Pearson (PCC) and Spearman coefficients. Gene-content similarity varies widely even among pairs with near-identical ANI, as quantified in (B). **(B)** Distribution of GCSj values across binned ANI ranges. Violin plots show the density distribution of GCSj, with internal boxplots indicating the median and interquartile range (IQR). The number of pairwise comparisons (*n*) is shown above each bin. The variance of GCSj is not homogeneous across ANI ranges (Fligner-Killeen test, p-value < 2.2e-16), and the spread of gene-content similarity peaks at the highest sequence identities (ANI 99.5-100%), highlighting extensive accessory genome variation among closely related strains.

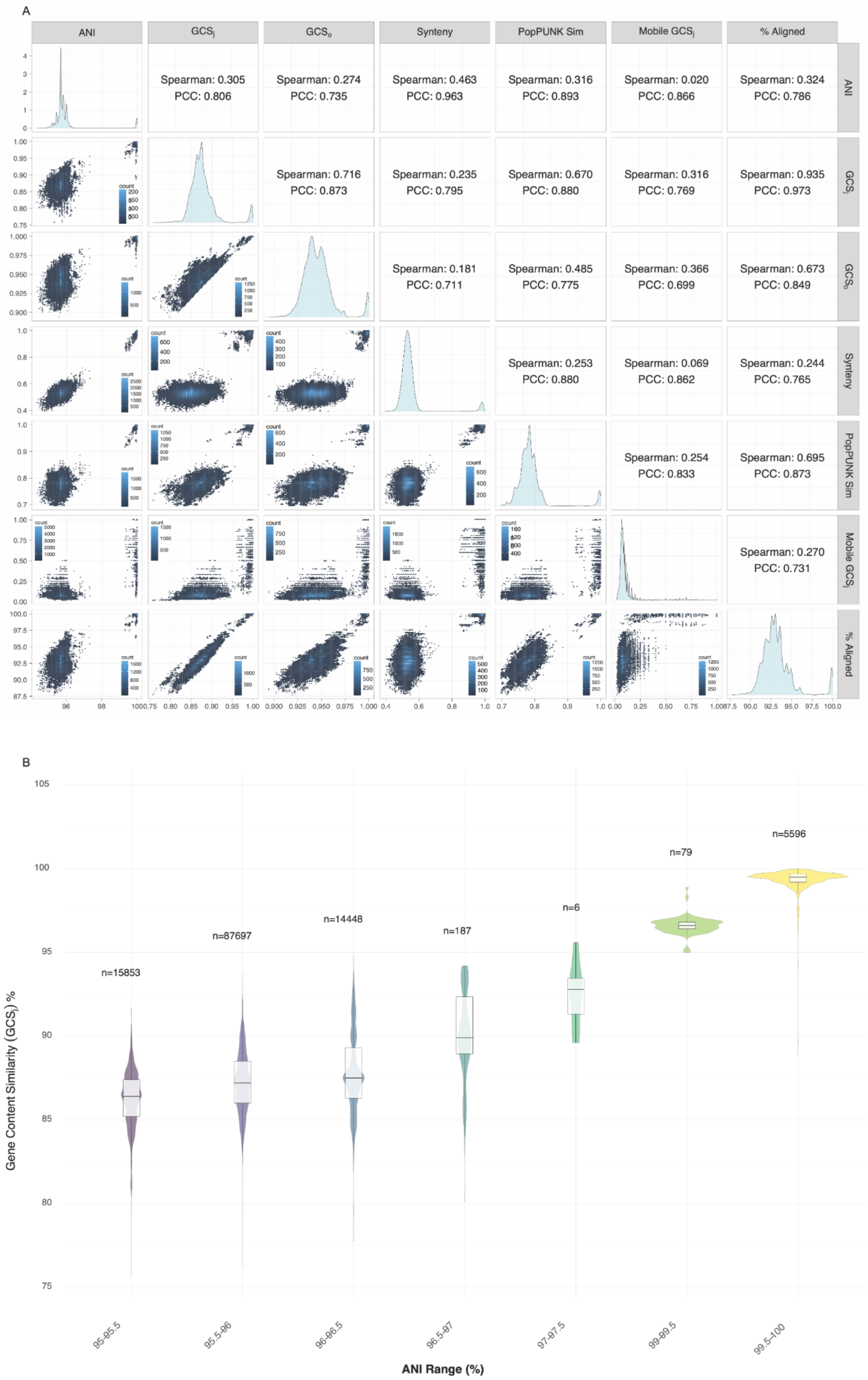


**Supplementary Figure S4. Comparing pairwise similarity measures for *H. pylori* genomes. (A)** Pairwise comparisons of similarity measures: ANI; gene-content similarity by Jaccard (GCSj) and overlap coefficient (GCSo); synteny score (SynTracker); PopPUNK accessory similarity; mobile-element profile similarity (Mobile GCSj); and fraction of aligned bases (% Aligned). Upper-right panels give Pearson (PCC) and Spearman coefficients. **(B)** Distribution of GCSj values across binned ANI ranges. Violin plots show the density distribution of GCSj, with internal boxplots indicating the median and IQR; n is shown above each bin. As expected given the limited accessory genome diversity of this species, GCSj variance remains low across all ANI ranges (Fligner-Killeen test, $p < 2.2e\text{-}16$, but IQR ≤ 3.4% throughout and only 0.5% at ANI > 99.5%), in contrast to the pattern observed in *E. coli*, *P. aeruginosa*, and *S. aureus*.

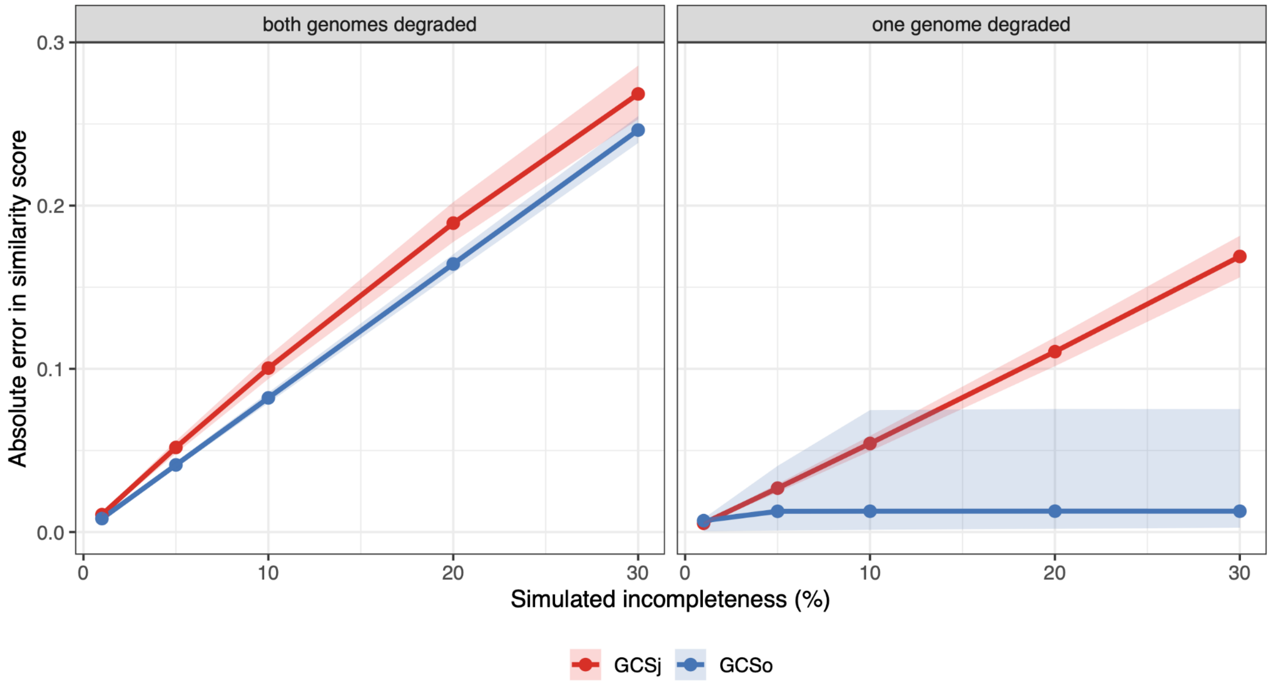


**Figure S5. GCSo is highly robust to simulated genome incompleteness.** Random incompleteness was simulated in the *E. coli* phyletic-pattern matrix by deleting a fraction of present genes from either one or both genomes in randomly selected genome pairs. For each degraded pair, GCSj (red) and GCSo (blue) were recalculated and compared against the baseline similarity computed from the complete genomes. Solid lines indicate the median absolute error across simulated pairs, and shaded ribbons represent the interquartile range. In both scenarios, GCSo exhibited lower error than GCSj, with the most striking improvement observed when only one genome in a pair was degraded. Simulations were performed on 20,000 genome pairs, with 10 replicate draws per pair at each indicated incompleteness level (1%, 5%, 10%, 20%, and 30%).

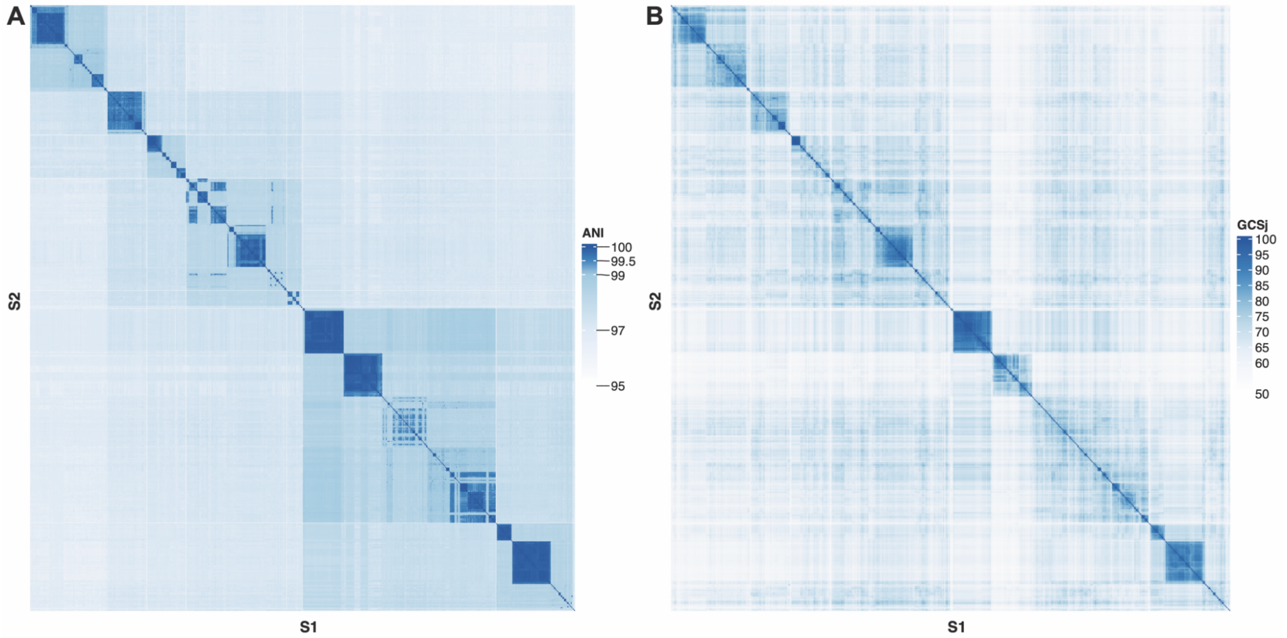


**Figure S6. Nested ANI-GCSj similarity matrices for *E. coli* genomes.** The ANI (**A**) and GCSj similarity (**B**) matrices for all pairwise comparisons of *E. coli* genomes. Genomes were first grouped into ten coarse ANI-based clusters using hierarchical clustering of ANI distances. Within each ANI-defined group, genomes were reordered according to GCSj similarity. The same genome order was used for both matrices. The GCSj matrix reveals additional gene-content variation within ANI-defined groups, visible as internal banding and substructure.

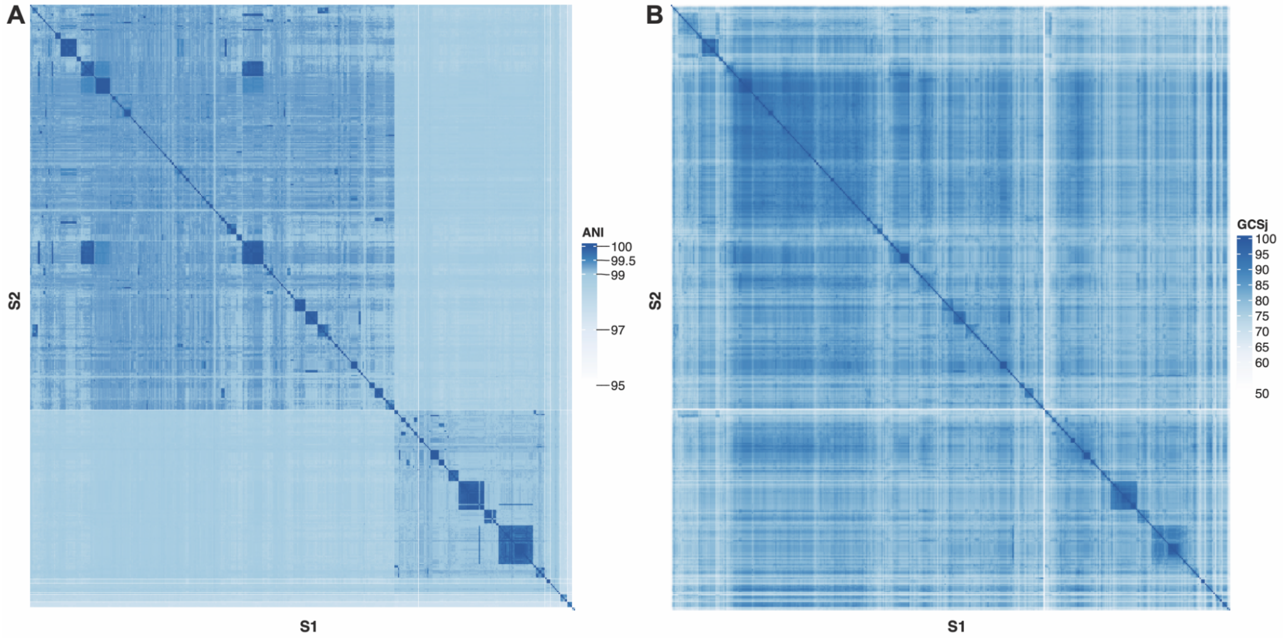


**Figure S7. Nested ANI-GCSj similarity matrices for *P. aeruginosa* genomes.** The ANI (**A**) and GCSj similarity (**B**) matrices for all pairwise comparisons of *P. aeruginosa* genomes. Genomes were first grouped into ten coarse ANI-based clusters using hierarchical clustering of ANI distances. Within each ANI-defined group, genomes were reordered according to GCSj similarity. The same genome order was used for both matrices. The GCSj matrix reveals additional gene-content variation within ANI-defined groups, visible as internal banding and substructure.

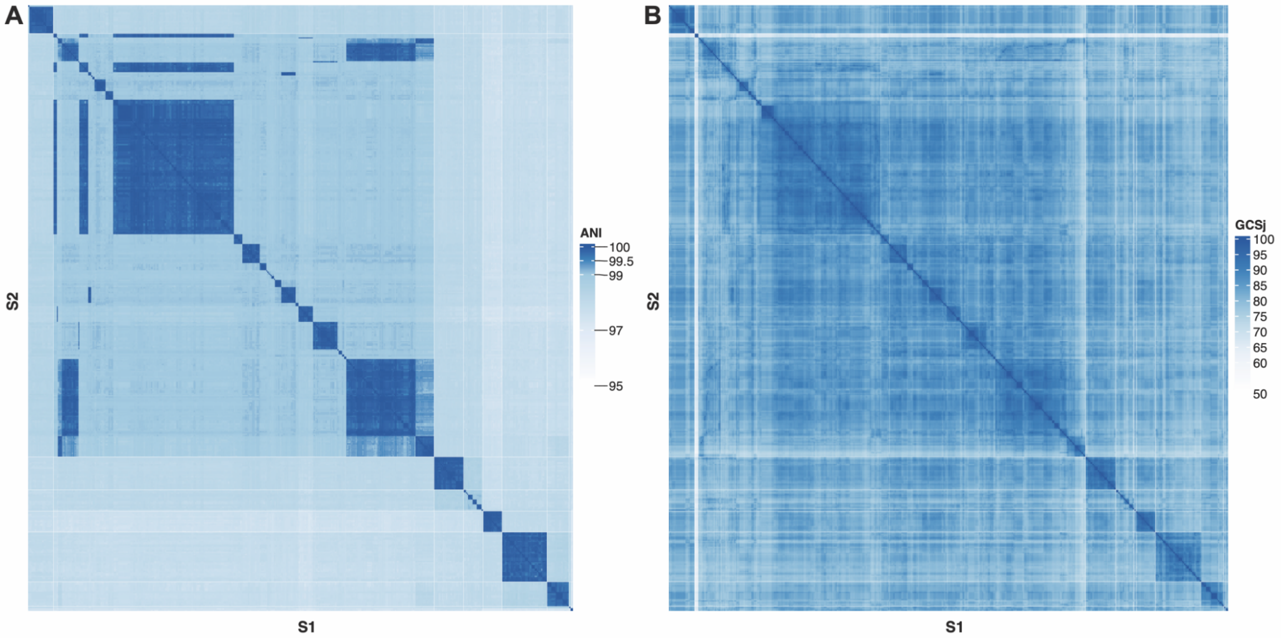


**Figure S8. Nested ANI-GCSj similarity matrices for *S. aureus* genomes.** The ANI (**A**) and GCSj similarity (**B**) matrices for all pairwise comparisons of *S. aureus* genomes. Genomes were first grouped into ten coarse ANI-based clusters using hierarchical clustering of ANI distances. Within each ANI-defined group, genomes were reordered according to GCSj similarity. The same genome order was used for both matrices. The GCSj matrix reveals additional gene-content variation within ANI-defined groups, visible as internal banding and substructure.

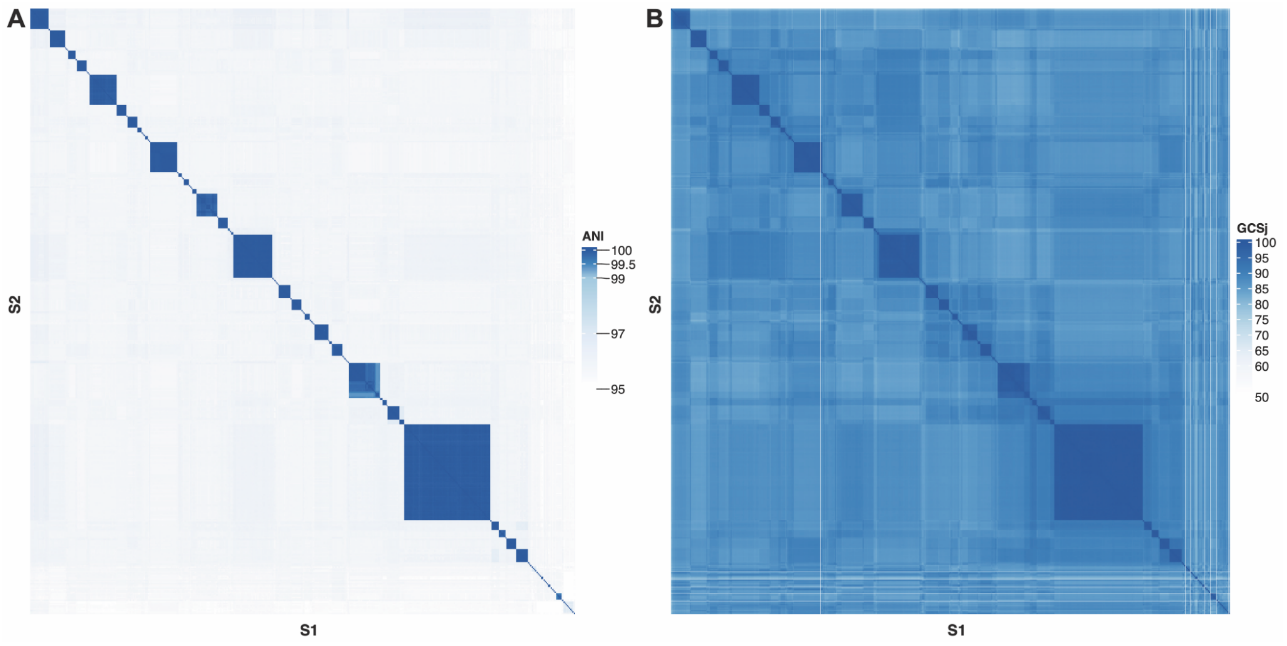


**Figure S9. Nested ANI-GCSj similarity matrices for *H. pylori* genomes.** The ANI (**A**) and GCSj similarity (**B**) matrices for all pairwise comparisons of *H. pylori* genomes. Genomes were first grouped into ten coarse ANI-based clusters using hierarchical clustering of ANI distances. Within each ANI-defined group, genomes were reordered according to GCSj similarity. The same genome order was used for both matrices. Compared with the other species, *H. pylori* shows less pronounced GCSj heterogeneity, consistent with lower gene-content variability.

**A.**

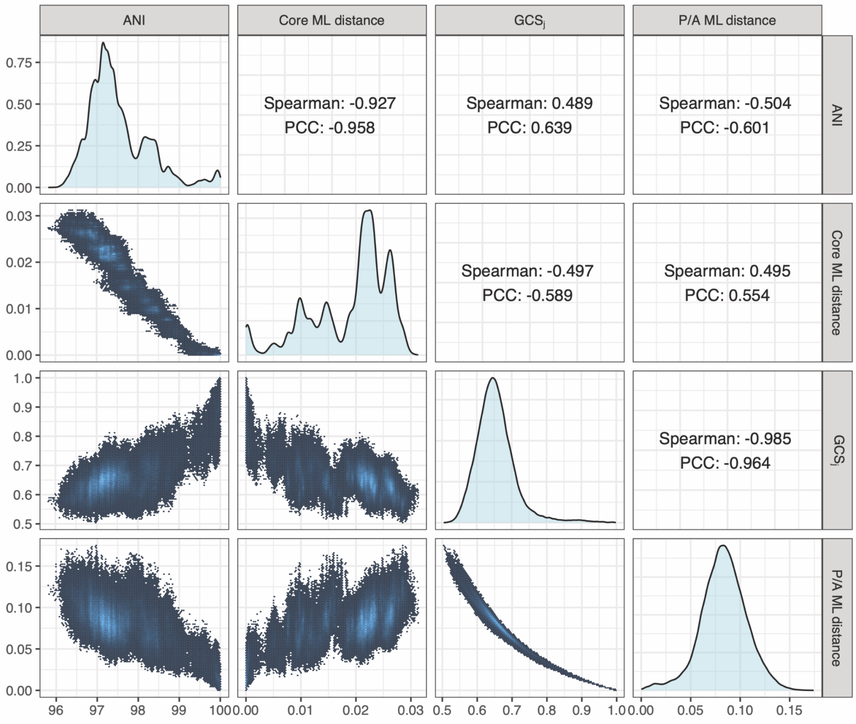
ANI
Core ML distance
$GCS_j$
P/A ML distance
Spearman: -0.927
PCC: -0.958
Spearman: 0.489
PCC: 0.639
Spearman: -0.504
PCC: -0.601
Spearman: -0.497
PCC: -0.589
Spearman: 0.495
PCC: 0.554
Spearman: -0.985
PCC: -0.964

**B.**

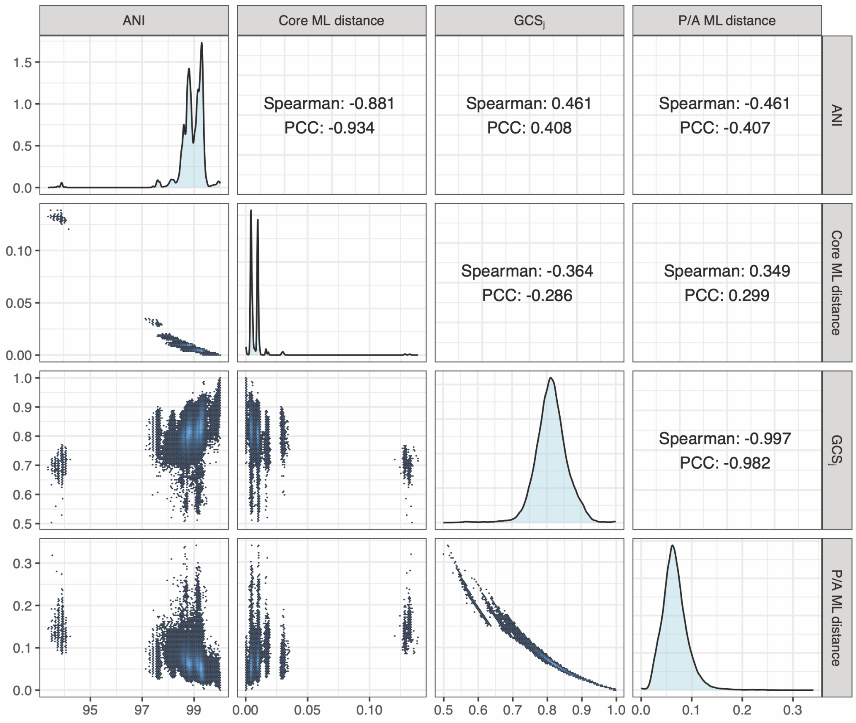
ANI
Core ML distance
$GCS_j$
P/A ML distance
Spearman: -0.881
PCC: -0.934
Spearman: 0.461
PCC: 0.408
Spearman: -0.461
PCC: -0.407
Spearman: -0.364
PCC: -0.286
Spearman: 0.349
PCC: 0.299
Spearman: -0.997
PCC: -0.982

**C.**

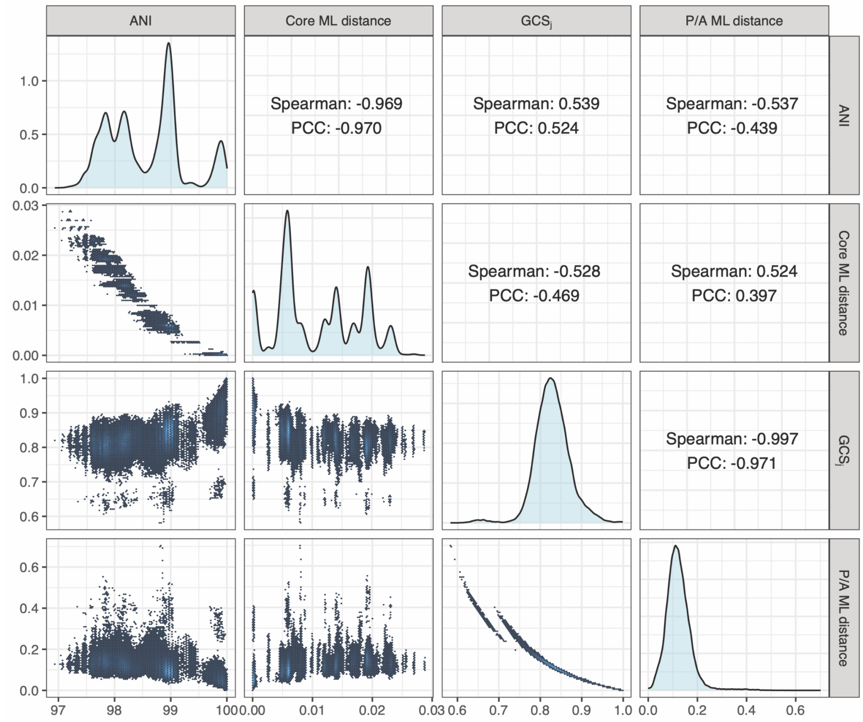


**D.**

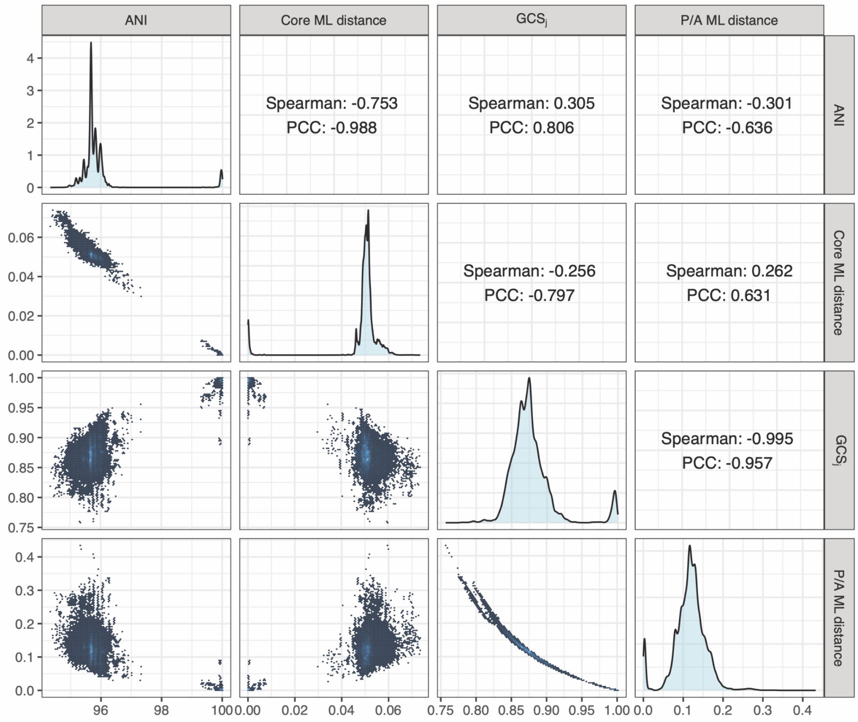


**Figure S10: Comparing ANI and GCSj with maximum likelihood (ML) distances based on nucleotides-core-genome and gene-content phyletic-pattern.** (**A**) *E. coli*, (**B**) *P. aeruginosa*, (**C**) *S. aureus*, and (**D**) *H. pylori*.

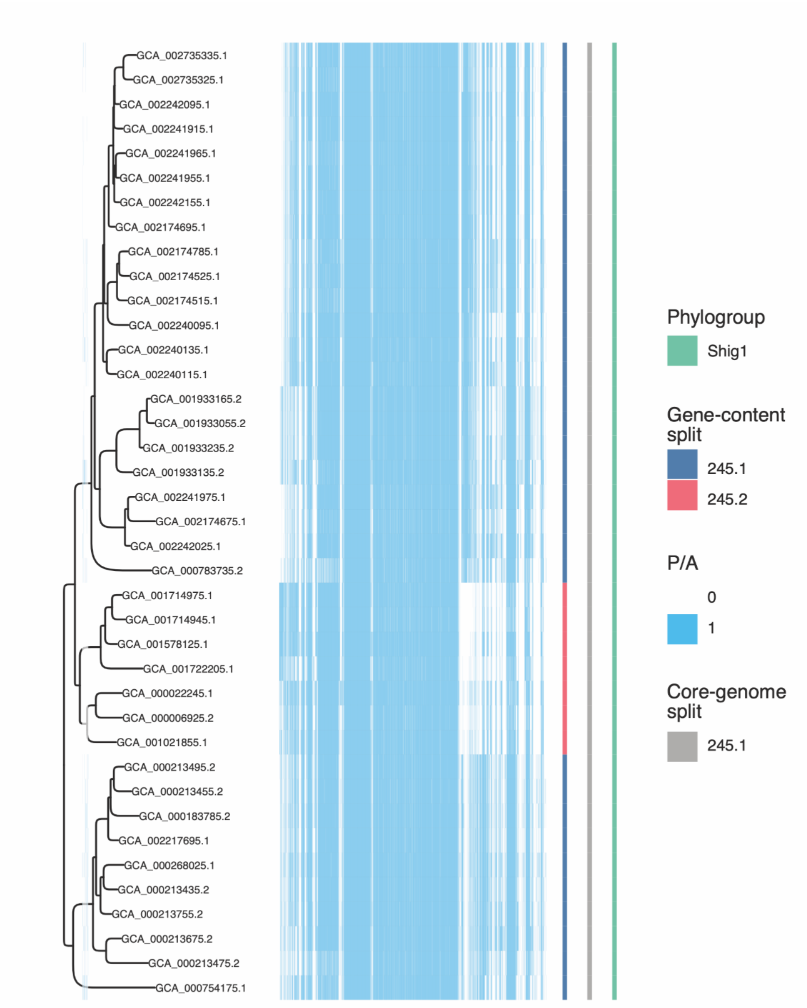


**Figure S11. Gene-content-based tree reveals additional subdivision within *E. coli* ST245.** Zoomed view of the gene-content-based maximum-likelihood tree for ST245 isolates (see Figure 2A), with the corresponding orthology-group presence/absence matrix (blue = present, white = absent) shown alongside. Colored tracks indicate phylogroup assignment and clade assignments from the gene-content-based and core-genome-based trees. All ST245 isolates remain a single, undivided group in the core-genome tree, whereas the gene-content-based split resolves them into two distinct groups (245.1 and 245.2), indicating that this subdivision is detectable through gene content but not through core-genome sequence similarity alone.

.

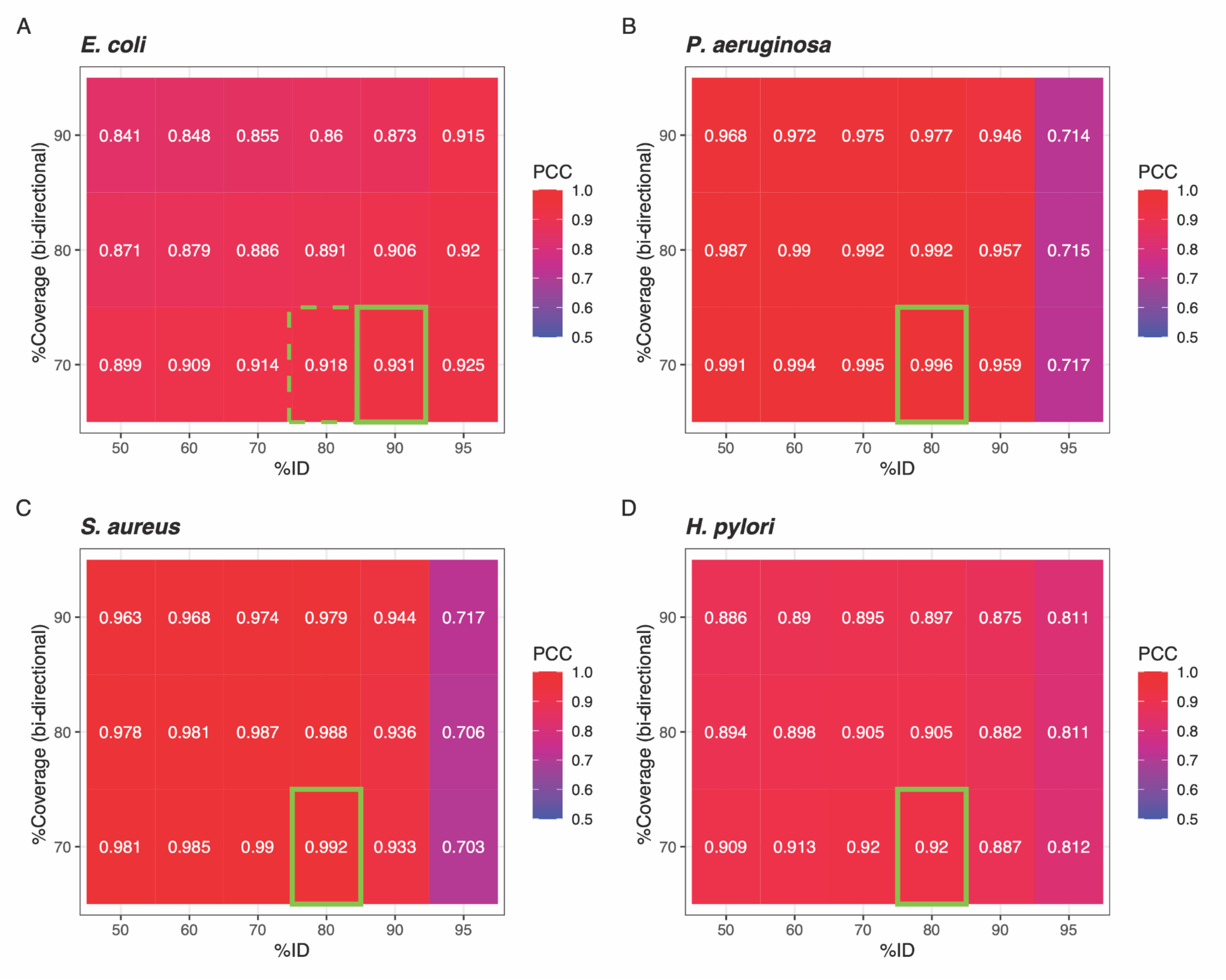


**Figure S12. Pearson correlation coefficient (PCC) between PanX-based GCSj and DIAMOND-DeepClust-based GCSj across combinations of percent identity and bi-directional coverage thresholds.** (**A**) *E. coli*, (**B**) *P. aeruginosa*, (**C**) *S. aureus*, (**D**) *H. pylori*. For each species, the solid green rectangle marks the combination yielding the highest PCC. A fixed threshold of 70% bidirectional coverage / 80% identity performs best in *P. aeruginosa*, *S. aureus*, and *H. pylori*, and performs comparably well in E. coli (dashed green rectangle; PCC = 0.918, the fourth-highest value, versus PCC = 0.931 for the best combination). This indicates that GCSj values are overall robust to the choice of clustering thresholds across a broad range of identity/coverage thresholds, supporting the use of a single fixed threshold across species.

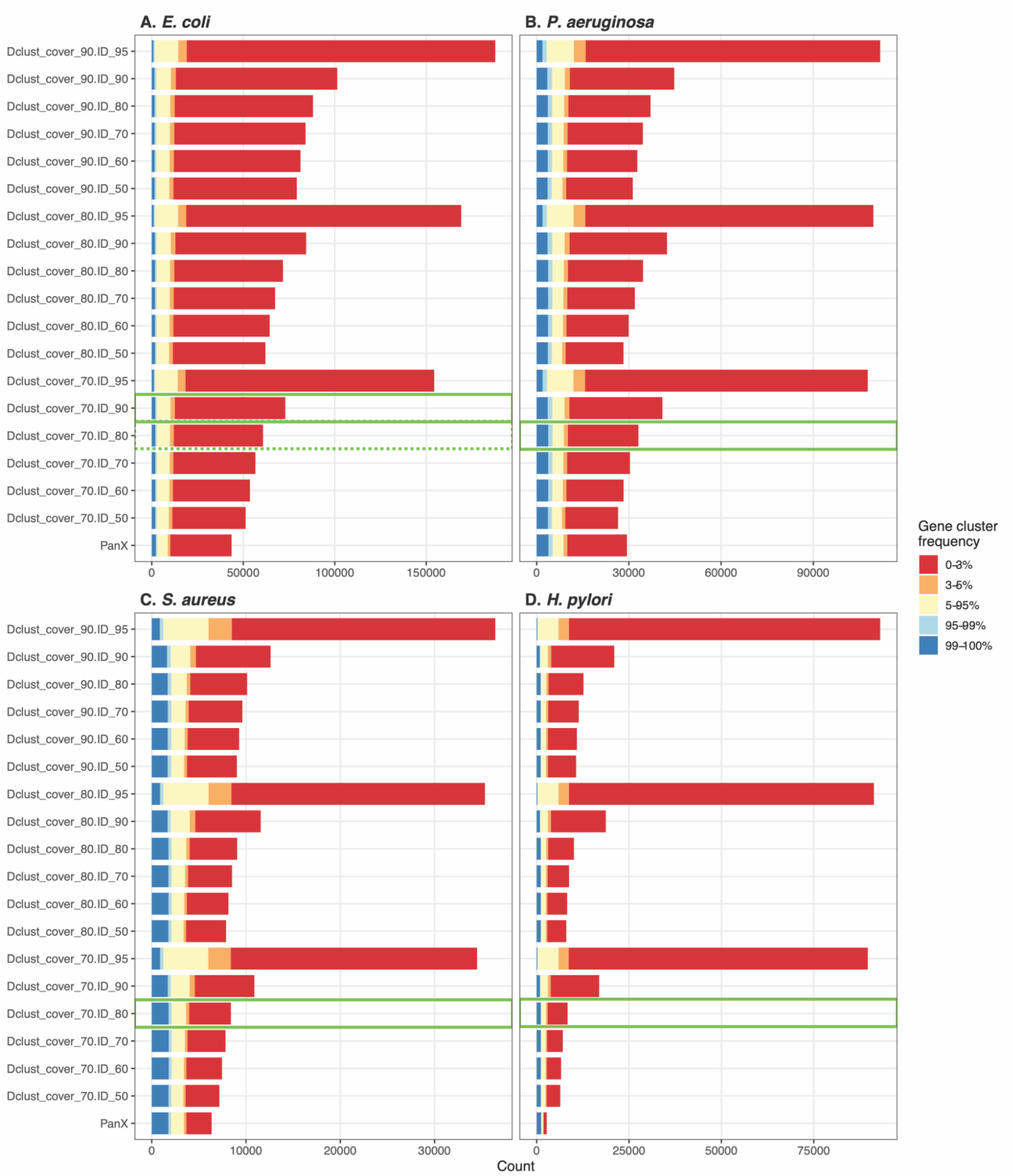


**Figure S13. Comparison of gene-cluster frequency-class composition between PanX orthology groups and DIAMOND-DeepClust clusters across combinations of percent identity and bi-directional coverage.** For each clustering approach, gene clusters were binned by the fraction of genomes in which they occur (99-100%, 95-99%, 5-95%, 3-5%, 0-3%), and the number of clusters in each frequency class is shown as a stacked bar. Combinations with high GCSj Pearson correlation coefficient (PCC) to PanX (see Figure S12) are indicated by green rectangles. (**A**) *E. coli*, (**B**) *P. aeruginosa*, (**C**) *S. aureus*, and (**D**) *H. pylori*. DIAMOND-DeepClust at higher identity thresholds inflates the rare-cluster (0-3%) count by over-splitting, whereas the recommended 80/70 threshold yields a frequency-class composition close to PanX.

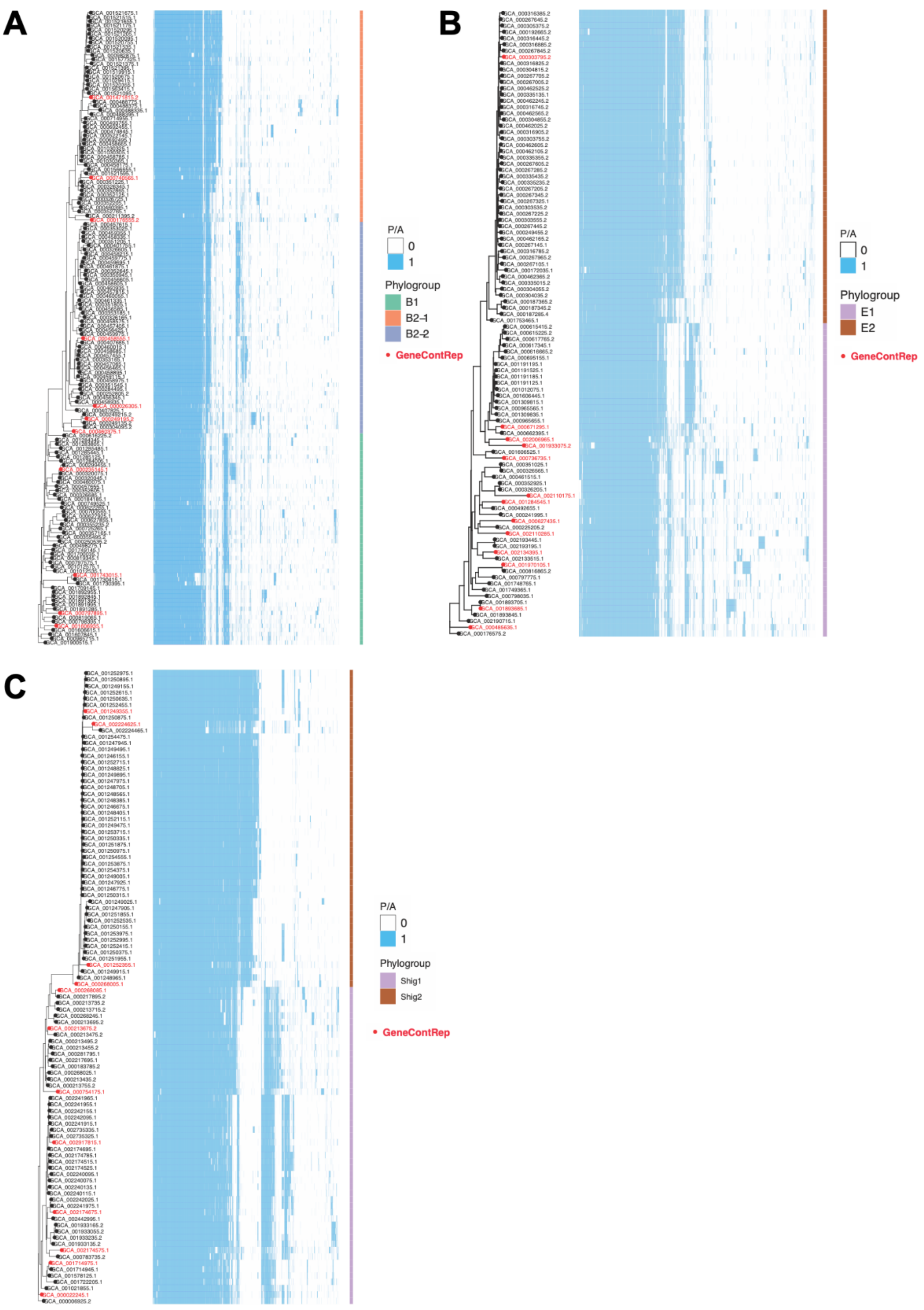


**Figure S14. Representative genomes selected using GeneContRep.** (**A**) *E. coli* phylogroup B: 11 representative genomes were selected using a GCSj cutoff of 0.7. (**B**) *E. coli* phylogroup E: 13 representative genomes were selected using a GCSj cutoff of 0.7. (**C**) Shigella: 12 representative genomes were selected using a GCSj cutoff of 0.8 (a slightly higher cutoff was used for Shigella to account for its lower gene-content diversity while achieving a comparable number of representatives). Presence/absence profiles of PanX orthology groups (blue) are plotted for each genome, indicating the presence (blue) or absence (white) of specific genes, ordered according to the maximum likelihood tree inferred with IQ-TREE. Orthology groups present in only one genome were not plotted. Colored side tracks indicate phylogroup/sub-phylogroup assignment (B1, B2-1, B2-2 in A; E1, E2 in B; Shig1, Shig2 in C), following the MASH-based subdivisions of Abram et al. (2021). Representative genomes selected by GeneContRep are marked in red.

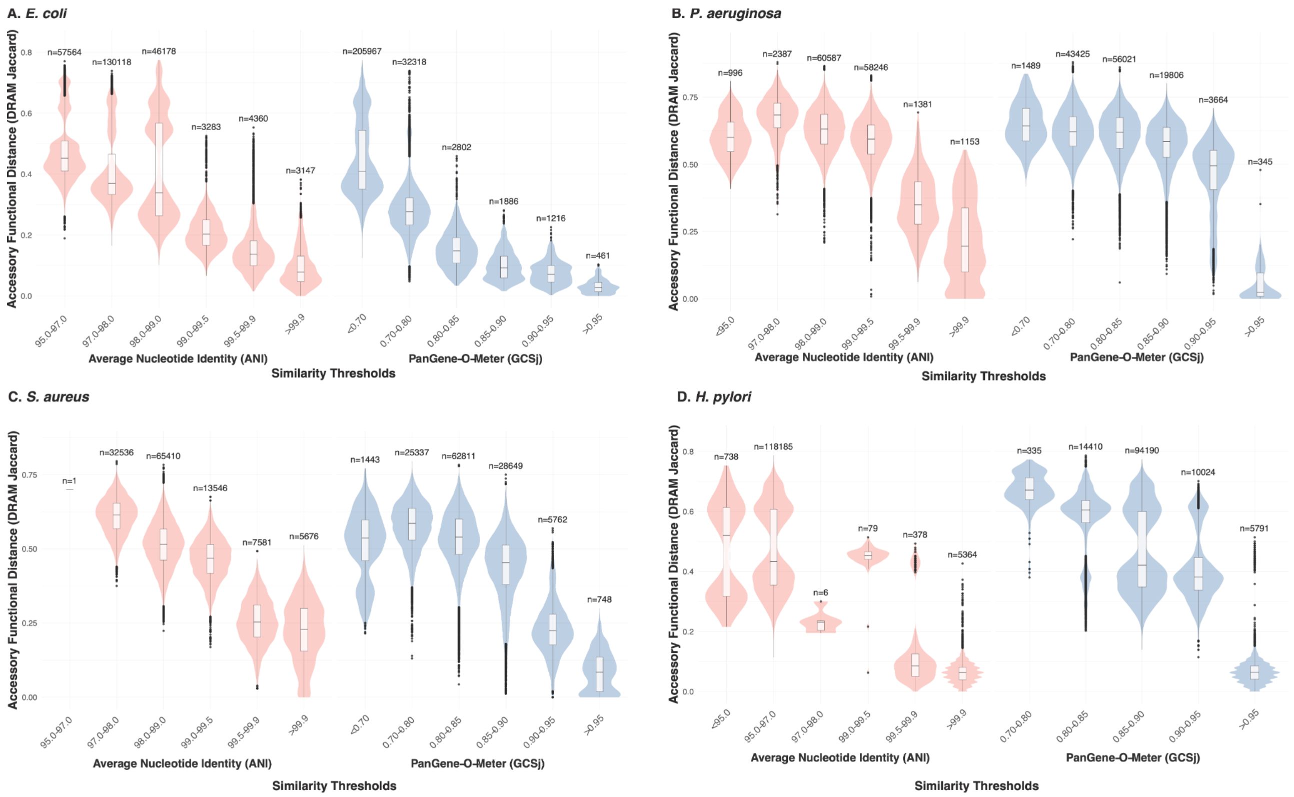


**Figure S15. GCSj captures functional diversity unresolved by nucleotide identity across four bacterial species.** Pairwise accessory functional distances (Jaccard distance of DRAM-derived KEGG functional profiles) are shown for genome pairs grouped by ANI (pink, left panels) or GCSj (blue, right panels) similarity thresholds. Numbers above each violin indicate the number of genome pairs per bin; note that some bins, particularly at the extremes, contain very few pairs (e.g., H. pylori ANI 97.0-98.0%, n = 6) and should be interpreted with caution. Boxes show median and IQR; whiskers extend to 1.5×IQR. Results are shown for (**A**) *Escherichia coli*, (**B**) *Pseudomonas aeruginosa*, (**C**) *Staphylococcus aureus*, and (**D**) *Helicobacter pylori*. Across all species, genome pairs remained functionally diverse even within the highest ANI bins, whereas functional distance decreased sharply and consistently with increasing GCSj, converging to its lowest values in the highest GCSj bin (>0.95) in every species. This pattern held even in H. pylori, whose comparatively closed pan-genome and limited accessory gene diversity yield a smaller absolute gap between ANI- and GCSj-based resolution than in the other three species, but the same qualitative trend.

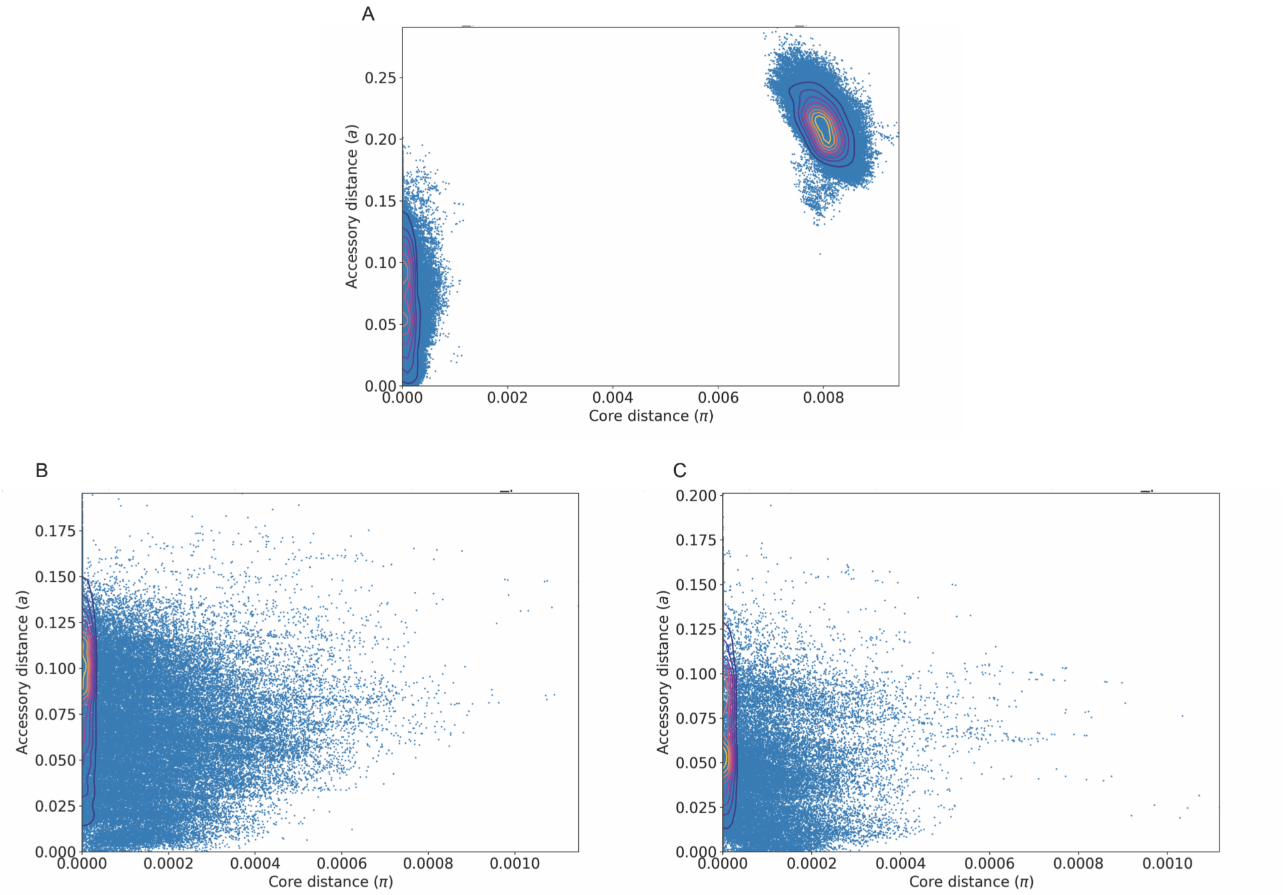


**Figure S16. PopPUNK distance distribution for Klebsiella pneumoniae ST258 and ST307.** Each point represents a pairwise genome comparison. Core distance (π) reflects nucleotide divergence in shared sequences; accessory distance (a) reflects k-mer-based estimation of accessory genome divergence; both were calculated by PopPUNK. (**A**) Combined view of all pairwise comparisons for both STs. Between-ST comparisons (right cluster, core distance ≈ 0.008) are clearly separated from within-ST comparisons (left cluster, core distance ≈ 0), confirming the distinct evolutionary trajectories of the two lineages. (**B, C**) The same within-ST comparisons from the left cluster in (A), shown separately for ST258 (**B**) and ST307 (**C**), with core distances rescaled to the 0-0.001 range. In both lineages, within-ST accessory distances span a broad, continuous range with no resolvable substructure, in contrast to the fine-grained resolution achieved by GCSj (see main text; Figures 5, 6).

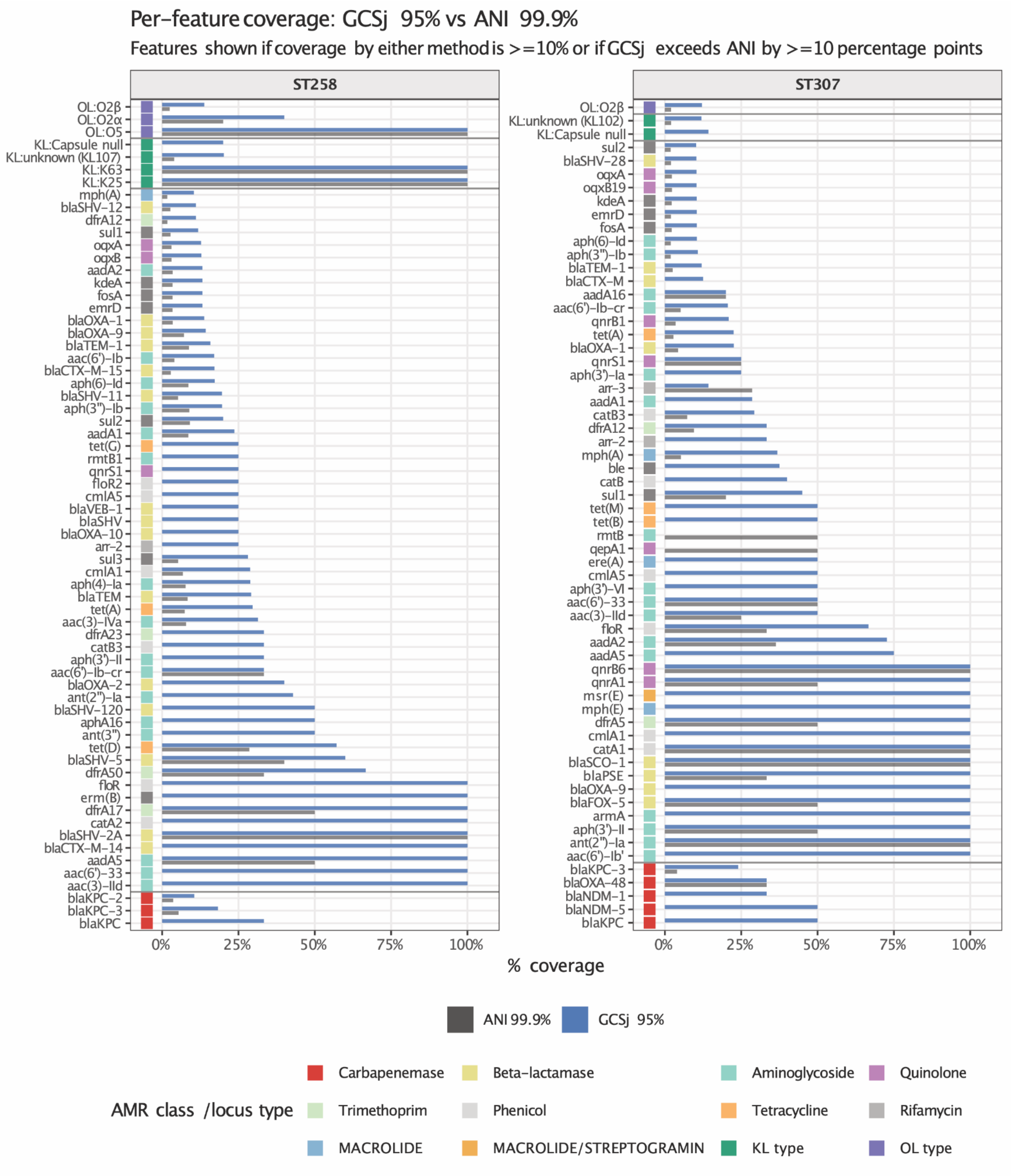


**Figure S17. Comprehensive per-feature coverage comparison for ANI and GCSj representatives in *Klebsiella pneumoniae*.** Bars show the fraction of carriers retained by ANI 99.9% representatives (grey) or GCSj 95% representatives (blue), relative to the total number of carriers in the full ST258 or ST307 cohort (500 genomes each). Features are shown if either method retained at least 10% of carriers or if GCSj exceeded ANI by at least 10 percentage points. Colored squares indicate AMR class or locus type. Results are shown separately for ST258 (left) and ST307 (right). Across both lineages, GCSj 95% representatives consistently achieve higher coverage than ANI 99.9% representatives across all represented AMR classes, including carbapenemases (red; e.g., blaKPC, blaNDM-1), beta-lactamases (yellow), aminoglycosides (teal), macrolides (light blue), quinolones (purple), tetracyclines (dark teal), trimethoprim (light green), phenicols (grey-green), and macrolide/streptogramin determinants (orange), as well as capsule KL types and O-locus OL types. The most striking differences are observed for carbapenemase genes, which are largely absent from ANI 99.9% representatives but substantially recovered by GCSj 95% representatives in both lineages.

**Table S1**

| Species | Measure | Min. | 1st Qu. | Median | Mean | 3rd Qu. | Max. | SD |
|---|---|---|---|---|---|---|---|---|
| ***Escherichia coli*** | Genome length (Mbp) | 4.225 | 4.980 | 5.194 | 5.159 | 5.368 | 6.323 | 0.304 |
| | Number of genes | 3,845 | 4,721.75 | 4,963.5 | 4,976.133 | 5202 | 6111 | 369.558 |
| | N50 (Mbp) | 0.019 | 0.094 | 0.150 | 1.141 | 1.325 | 5.697 | 1.731 |
| | Number of contigs | 1 | 14 | 122.5 | 156.84 | 244 | 806 | 154.312 |
| | Completeness (%) | 99.5 | 100 | 100 | 99.997 | 100 | 100 | 0.027 |
| | Contamination (%) | 0 | 0.09 | 0.25 | 0.468 | 0.63 | 4.83 | 0.604 |
| | GC content | 50 | 50 | 51 | 50.706 | 51 | 51 | 0.456 |
| | Number of STs | 193 | | | | | | |
| ***Pseudomonas aeruginosa*** | Genome length (Mbp) | 6.015 | 6.488 | 6.796 | 6.768 | 7.009 | 7.730 | 0.33 |
| | Number of genes | 3,900 | 5,891.2 | 6,207 | 6,178.3 | 6,429 | 7,289 | 373.423 |
| | N50 (Mbp) | 6.015 | 6.472 | 6.739 | 6.719 | 6.937 | 7.564 | 0.293 |
| | Number of contigs | 1 | 1 | 1 | 1.396 | 2 | 6 | 0.808 |
| | Completeness (%) | 95.51 | 100 | 100 | 99.975 | 100 | 100 | 0.303 |
| | Contamination (%) | 0.02 | 0.16 | 0.26 | 0.373 | 0.46 | 2.24 | 0.334 |
| | GC content | 65 | 66 | 66 | 66.028 | 66 | 67 | 0.322 |
| | Number of STs | 186 | | | | | | |
| ***Staphylococcus aureus*** | Genome length (Mbp) | 2.677 | 2.776 | 2.827 | 2.832 | 2.879 | 3.132 | 0.072 |
| | Number of genes | 1,986 | 2,594 | 2,649.5 | 2,657.6 | 2,719 | 3,102 | 105.094 |
| | N50 (Mbp) | 2.677 | 2.761 | 2.807 | 2.815 | 2.859 | 3.089 | 0.068 |
| | Number of contigs | 1 | 1 | 2 | 1.804 | 2 | 6 | 0.885 |
| | Completeness (%) | 100 | 100 | 100 | 100 | 100 | 100 | 0 |
| | Contamination (%) | 0 | 0.08 | 0.12 | 0.201 | 0.21 | 3.28 | 0.293 |
| | GC content | 33 | 33 | 33 | 33 | 33 | 33 | 0 |
| | Number of STs | 87 | | | | | | |
| ***Helicobacter pylori*** | Genome length (Mbp) | 1.538 | 1.595 | 1.637 | 1.628 | 1.660 | 1.717 | 0.039 |
| | Number of genes | 1,425 | 1,504 | 1,536 | 1,532 | 1,563.2 | 1,623 | 36.682 |
| | N50 (Mbp) | 0.006 | 0.064 | 0.08 | 0.084 | 0.097 | 0.230 | 0.033 |
| | Number of contigs | 26 | 44 | 54 | 61.894 | 68 | 460 | 37.8 |
| | Completeness (%) | 99.01 | 99.99 | 99.99 | 99.988 | 99.99 | 100 | 0.046 |
| | Contamination (%) | 0 | 0.01 | 0.07 | 0.205 | 0.22 | 4.94 | 0.440 |
| | GC content | 39 | 39 | 39 | 39 | 39 | 39 | 0 |
| | Number of STs | 21 | | | | | | |

**Table S2 - Assembly IDs for the genomes analyzed in this study**

Table S2 is available online

**Table S3 - DIAMOND-DeepClust reduces gene-content clustering runtime from hours to minutes compared with PanX orthology assignment.**

| Species | Genomes | PanX (steps 03–06) | PanX threads | step06 share | DIAMOND-DeepClust (64 threads) | Speedup |
|---|---|---|---|---|---|---|
| *E. coli* | 700 | 8 h 27 m | 128 | 82% | 2 m 46 s | ~183× |
| *P. aeruginosa* | 500 | 11 h 09 m | 96 | 85% | 2 m 12 s | ~303× |
| *S. aureus* | 500 | 2 h 09 m | 96 | 49% | 0 m 58 s | ~134× |
| *H. pylori* | 500 | 1 h 51 m | 64 | 63% | 0 m 47 s | ~143× |

* PanX runtimes are summed over pipeline steps 03-06 from the original analysis logs, at the thread counts indicated. DIAMOND-DeepClust (cluster + recluster) was re-run for all datasets on a single system at 64 threads, equal to (*H. pylori*) or fewer than (all other datasets) the threads available to PanX, making the comparison conservative. Runtimes are wall-clock and illustrative rather than a controlled benchmark. PanX additionally produces multiple sequence alignments and gene trees, which are not required for GCS calculation but were used for the core-genome phylogenetic analyses presented here.
